# Phospholipid isotope tracing reveals β-catenin-driven suppression of phosphatidylcholine metabolism in hepatocellular carcinoma

**DOI:** 10.1101/2023.10.12.562134

**Authors:** Chad VanSant-Webb, Hayden K. Low, Junko Kuramoto, Claire E. Stanley, Hantao Qiang, Audrey Su, Alexis N. Ross, Chad G. Cooper, James E. Cox, Scott A. Summers, Kimberley J. Evason, Gregory S. Ducker

## Abstract

**Background and Aims:** Activating mutations in the *CTNNB1* gene encoding β-catenin are among the most frequently observed oncogenic alterations in hepatocellular carcinoma (HCC). HCC with *CTNNB1* mutations show profound alterations in lipid metabolism including increases in fatty acid oxidation and transformation of the phospholipidome, but it is unclear how these changes arise and whether they contribute to the oncogenic program in HCC.

**Methods:** We employed untargeted lipidomics and targeted isotope tracing to quantify phospholipid production fluxes in an inducible human liver cell line expressing mutant β-catenin, as well as in transgenic zebrafish with activated β-catenin-driven HCC.

**Results:** In both models, activated β-catenin expression was associated with large changes in the lipidome including conserved increases in acylcarnitines and ceramides and decreases in triglycerides. Lipid flux analysis in human cells revealed a large reduction in phosphatidylcholine (PC) production rates as assayed by choline tracer incorporation. We developed isotope tracing lipid flux analysis for zebrafish and observed similar reductions in phosphatidylcholine synthesis flux accomplished by sex-specific mechanisms.

**Conclusions:** The integration of isotope tracing with lipid abundances highlights specific lipid class transformations downstream of β-catenin signaling in HCC and suggests future HCC-specific lipid metabolic targets.

**Synopsis:** In this work, we show by lipid specific isotope tracing that mutations in the oncogene *CTNNB1* leads to conserved changes in lipid metabolism in hepatocellular carcinoma. These include the stimulation of fatty acid oxidation and a suppression of phosphorylcholine synthesis.

## Introduction

The incidence of hepatocellular carcinoma (HCC) has tripled over the last four decades while the 5-year survival rates remain among the worst of all cancers (American Cancer Society, n.d.). Compared to other common solid tumor malignancies, HCC has unique features that have hindered basic biological understanding and made therapeutic advances difficult in this disease(Tovoli et al., 2018). These include a low frequency of targetable genetic mutations, a heterogenic metabolic profile, and a generally suppressed immune microenvironment(Pfister et al., 2021; Sangineto et al., 2020). Coupled with the natural history of the disease, these features have made HCC difficult to model in animal systems(Krutsenko et al., 2021; Nakagawa, 2015). Zebrafish (*Danio rerio*) are a vertebrate model organism with a complete immune system and metabolic network that show high conservation with mammals and have been used to study multiple aspects of tumor biology. The use of zebrafish tumor models has recently expanded to include the study of tumor metabolism and several groups have successfully exploited zebrafish to study the metabolism of melanoma (Lumaquin-Yin et al., 2023; Naser et al., 2021). We previously reported a zebrafish model of HCC driven solely by a *CTNNB1* transgene that recapitulates human HCC morphologically and transcriptionally in as little as 10 weeks (Evason et al., 2015).

Recurrent patterns of metabolic alterations support proliferative growth in cancer. The best characterized of these is the Warburg effect, whereby many tumors “overconsume” glucose even in the presence of oxygen, providing the mechanistic basis of ^18^FDG PET imaging of many solid tumors (Vander Heiden et al., 2009). However, enhanced glycolytic flux is only present in a minority of HCC cases, and PET imaging is not standard of care. Changes in lipid metabolism are just as frequent and are now appreciated to play a diverse role in mediating tumor cell biology (Broadfield et al., 2021). Tumor-specific alterations in fatty acid synthesis and oxidation help provide substrates for membrane construction and energy (Röhrig and Schulze, 2016), while changes in fatty acid saturation affect membrane fluidity and decrease tumor susceptibility to ferroptosis (Friedmann Angeli et al., 2019). Tumor cells make an array of signaling lipids such as prostaglandins and cholesterol-derived metabolites that interact with other cells in the tumor microenvironment including those of the immune system to evade detection and enhance proliferation (Zelenay et al., 2015). These discoveries have fueled interest in targeting various aspects of lipid metabolism, but the mechanisms giving rise to these changes and how critical they are for tumor growth are much less understood, limiting our ability to target tumor lipid biology therapeutically.

Mutations leading to the stabilization of the Wnt pathway transcriptional transactivator β-catenin (encoded by *CTNNB1*) are one of the most frequent oncogenic events in HCC, occurring in approximately 30% of tumors and defining a genetic subset of HCC (Ally et al., 2017; Calderaro et al., 2017; Nault et al., 2020). Preclinical and early clinical data suggest that *CTNNB1* mutations mediate some of the therapeutic intractability of HCC (Ruiz De Galarreta et al., 2019). Recent work demonstrated that activated β-catenin promotes fatty acid oxidation in osteoblasts and in an APC deletion-driven mouse model of HCC (Frey et al., 2018; Senni et al., 2019). β-catenin is a known regulator of several master lipid transcription factors including SREBP1 (Bagchi et al., 2020b, 2020a; Behari et al., 2010) but the complete effects of β-catenin on regulating lipid metabolism are unknown.

Changes in the class composition of the human HCC lipidome are widespread and have been measured by many groups using untargeted mass spectrometry analyses (Ismail et al., 2020; Krautbauer et al., 2016; Lu et al., 2018; Morita et al., 2013). However, many of the changes in lipid metabolism are inconsistent between studies likely reflecting the diverse etiologies of disease and mutational profiles among the patient populations sampled(Buechler and Aslanidis, 2020). Profiling of lipid metabolism by flux analysis and connecting metabolic and genetic phenotype may help to define pathways that are common (or unique) to different HCC etiologies and genetic subsets.

Metabolic flux, or the rate of production of a specific metabolite, provides information about the dynamics of a pathway, highlighting those that are most active. While metabolic flux cannot be directly observed, it can be calculated from turnover rates determined by stable isotope tracing analysis. The use of stable isotope tracers to quantify glucose and glutamine reaction fluxes *in vivo* has been transformational for our understanding of genotype and therapy-specific tumor metabolism (Faubert et al., 2021; Jang et al., 2018; Pavlova et al., 2022). Currently, the only fully developed method in tumor lipid metabolism is the use of deuterium oxide (D_2_O) to quantify rates of fatty acid synthesis (Zhang et al., 2017), which are elevated in breast, HCC, and prostate cancers (Kinlaw et al., 2016; Röhrig and Schulze, 2016; Zhou et al., 2022).

We focused on characterizing the lipid metabolism of a defined molecular driver of HCC, activated β-catenin. Here we report and define dysregulated lipid metabolism driven by activated β-catenin in immortalized human hepatocytes and zebrafish β-catenin-driven HCC. We discovered that activated β-catenin induced large changes in the lipidome in both systems, including conserved increases in acylcarnitines and ceramides and decreases in triglycerides. Targeted isotope tracing revealed a significant reduction in phosphatidylcholine (PC) production rates in human hepatocytes and in zebrafish HCC, wherein this decrease was accomplished by different sex-specific mechanisms. The integration of isotope tracing with lipid abundances and transcriptomics analysis highlights specific lipid class transformations driven by activated β-catenin signaling in HCC.

## Results

### Activated β-catenin drives lipidome remodeling in human hepatocytes

We first sought to determine the direct effects of activated β-catenin on lipidome reprograming in cultured human hepatocytes. THLE-2 cells, derived from normal human hepatocytes by infection with SV40 T antigen (Pfeifer et al., 1993), were transfected with plasmids encoding the Tet-on-Advanced system enabling precise control of expression of the target gene of interest. *CTNNB1* constructs encoded either *WT (WT-CTNNB1*) or mutant *CTNNB1* (*4x-Ala-CTNNB1*), an allele containing mutations (S33A, S37A, T41A, S45A) that prevent the normal phosphorylation and degradation of β-catenin protein (Supplemental Fig 1A). *WT-CTNNB1* or *4x-Ala-CTNNB1* cells were grown in selection media with neomycin and puromycin, and expression of *CTNNB1* was induced by treatment with doxycycline (Fig 1A). Addition of doxycycline increased *CTNNB1* transcript levels approximately 6-fold in both *WT-CTNNB1* and *4x-Ala-CTNNB1* cells (Supplemental Figure 1B). Expression of β-catenin protein was significantly higher in *4x-Ala-CTNNB1* cells treated with doxycycline compared to *WT-CTNNB1* cells with the same treatment (Fig 1B), consistent with the generation of a degradation-resistant protein. Doxycycline treatment increased expression of *GLUL*, a known β-catenin target gene, in both *WT-CTNNB1* and *4x-Ala-CTNNB1* cells Fig 1c) (Sekine et al., 2006). Overexpression of activated β-catenin, but not WT β-catenin, increased anchorage-independent cell growth (Fig 1D), while growth on collagen-treated surfaces was not affected (Supplemental Figure 1C), consistent with prior work (Orford et al., 1999). Thus, *4x-Ala-CTNNB1* cells show doxycycline-dependent induction of activated β-catenin leading to functional changes in cell growth and behavior.

**Figure 1:**
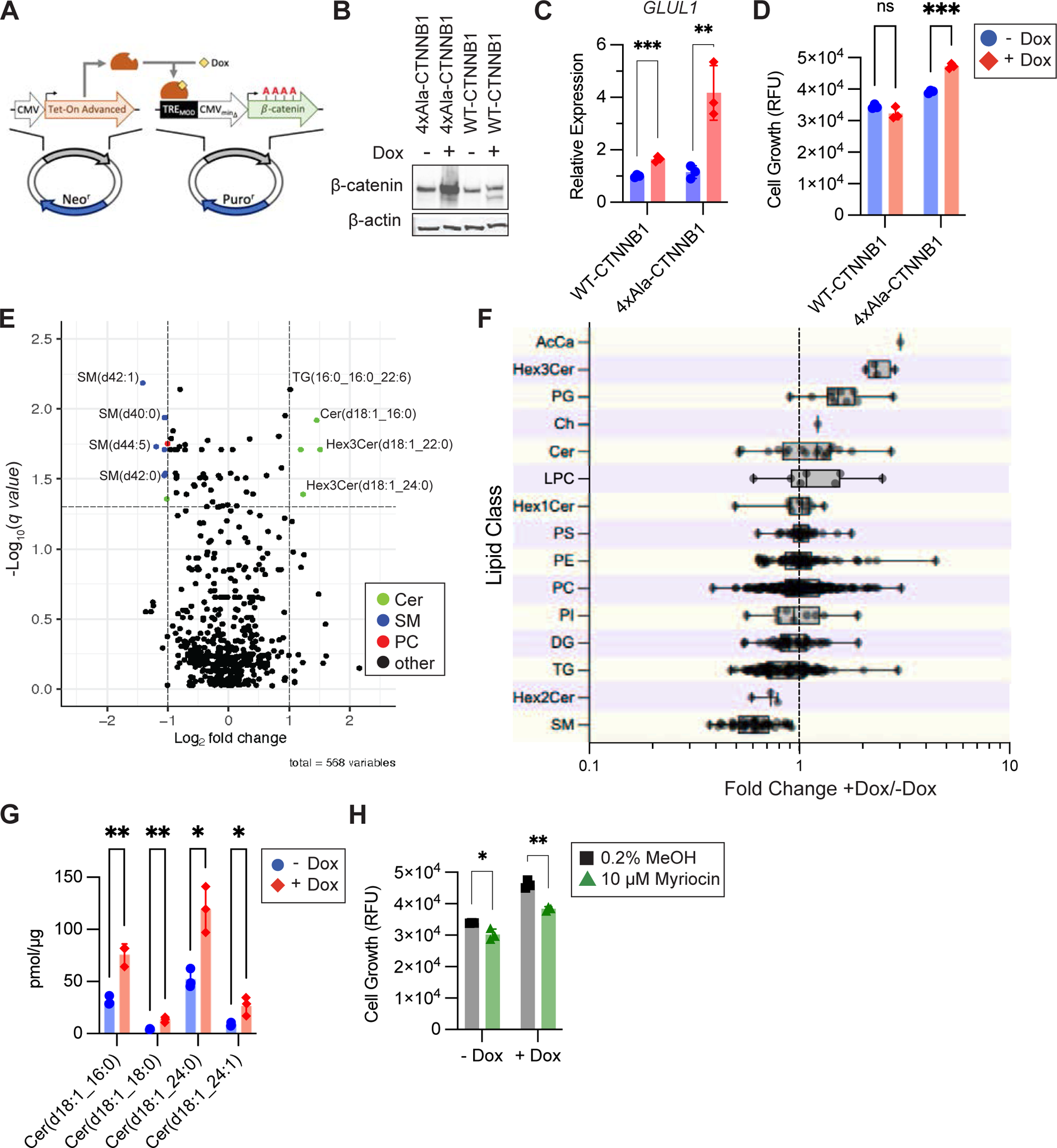
Mutant β-catenin expression drives lipidome remodeling in a human liver cell line. A. Overview of Tet-On Dox-inducible β-catenin system. B. Transformed but not tumorigenic THLE-2 cells were transfected with constructs encoding either mutant (4xAla) or WT *CTNNB1* +/− Dox induction and quantified by immunoblot. C. qRT-PCR of β-catenin-induced gene *GLUL* from THLE-2 cells +/− Dox treatment in *CTNNB1* overexpression constructs (n=3, mean±stdev). D. Anchorage independent growth assay of THLE-2 cells +/− Dox treatment in *CTNNB1* overexpression constructs (n=3, mean±stdev). E. Volcano plot showing change in abundance for individual lipid species from THLE-2 cells +/− Dox treatment in *4x-Ala-CTNNB1* overexpression construct (n=3). Y-axis is the −Log_10_q using a multiple-comparisons corrected false-discovery rate of 5%. Select individual species are named. F. Fold changes in lipid species by class. Lipid classes are AcCa: acylcarnitines, Hex3Cer: trihexosylceramides, PG: phosphatidylglycerol lipids, Ch: cholesterol, Cer: ceramides, LPC: lysophosphatidylcholine lipids, Hex1Cer: hexosylceramides, PS: phosphatidylserine lipids, PE: phosphatidylethanolamine lipids, PC: phosphatidylcholine lipids, PI: phosphatidylinositol lipids, DG: diacylglycerides, TG: triacylglycerides, Hex2Cer: dihexosylceramides, SM: sphingomyelins. Fold change is average change between *4x-Ala-CTNNB1* expressing THLE-2 cells +/− Dox (n=3). G. Normalized abundances of 18:1 ceramides in THLE-2 cells +/− Dox treatment in *4x-Ala-CTNNB1* overexpression construct (n=3, mean±stdev). H. Anchorage independent growth of THLE-2 cells +/− Dox treatment in *4x-Ala-CTNNB1* overexpression construct with or without serine palmitoyl transferase inhibitor myriocin (n=3, mean±stdev). RFU = relative fluorescence units. For C, D, G, and H, * p<.05, ** p<.01, *** p<.001, **** p<.0001 by multiple unpaired t-tests, corrected for multiple comparisons using the Holm-Šídák method.

To understand the metabolic changes driven by activated β-catenin expression, we used liquid chromatography-mass spectrometry (LC-MS) to quantify changes in the lipidome of *4x-Ala-CTNNB1* cells with and without doxycycline. Cells were cultured in adherent growth conditions and lipids were extracted before being identified and quantified using untargeted LC-MS. Using univariate analysis with a significance cut-off of at least 2-fold average change and a multiple-comparisons corrected false-discovery rate of 0.05, we identified 15 lipid species that were differentially expressed upon activated β-catenin expression (Fig 1E). With the one exception of triglyceride 16:0_16:0_22:6, all lipid species increased by doxycycline treatment were ceramides. Sphingomyelin (SM) lipids were the large majority of down-regulated lipids.

To highlight conserved changes that might occur among biochemically related molecules, we examined changes in lipid abundance grouped by class (Fig 1F). Grouping doxycycline-induced changes by lipid class revealed that trihexosylceramide (Hex3cer) lipids were upregulated in cells with activated β-catenin, as were most phosphatidylglycerol (PG) lipids (Fig 1F). SM lipids were universally downregulated, and the bulk of triglyceride species were also reduced upon β-catenin activation. Within other lipid classes, no clear trends were apparent. The absolute abundance of lipids organized by class was only different in doxycycline-treated cells for SM lipids (Supplemental Figure 1D).

The sole acyl carnitine species detected in our cell assay was significantly increased upon doxycycline induction. β-catenin induction also increased the expression of the rate-limiting enzyme in acyl-carnitine synthesis, *CPT1* (Supplemental Figure 1E). The increase in acyl-carnitine—the committed intermediary between fatty acids and their oxidation in mitochondria—together with the increase in *CPT* expression, suggests increased rates of fatty acid β-oxidation in hepatocytes expressing activated β-catenin.

To test the possibility that the lipidomic changes noted above could represent an artifact of doxycycline exposure, we compared the lipidomes of *WT-CTNNB1* cells with and without doxycycline treatment (Supplemental Figure 1F). Using the same FDR cut-off of 0.05, no individual lipid species were significantly altered upon doxycycline treatment. We observed an attenuated change in most lipid class averages, including PG, Hex3cer and triglyceride lipid classes, which no longer showed significant differences with and without doxycycline treatment (Supplemental Figure 1G). Although the individual detected lipids were not statistically significant, we observed a similar downregulation of SM lipids with doxycycline treatment between mutant and WT *CTNNB1-*expressing cells (Supplemental Figure 1G). These results confirm that most significant changes in lipid abundances observed in *4x-Ala-CTNNB1* cells upon doxycycline treatment are due to the presence of mutant β-catenin activity, rather than to the effects of doxycycline treatment or additional WT β-catenin. Further, the lack of significant changes in PG, Hex3cer, and triglyceride lipids with WT *CTNNB1* overexpression supports the hypothesis that the doxycycline-induced alterations in these lipid species in *4x-Ala-CTNNB1* cells are specifically related to overexpression of mutant *CTNNB1*.

Ceramides have potent roles in cellular lipid signaling and organismal physiology(Chaurasia and Summers, 2021). We observed that several ceramides comprising the canonical d18:1 sphingoid backbone were increased upon doxycycline treatment (Fig 1G). Ceramide levels are governed by a complex network of degrading pathways and synthesizing enzymes that can generate ceramides from other lipid classes or *de novo* from serine. We asked whether blocking the sole *de novo* synthesis pathway of ceramides with myriocin, an inhibitor of serine palmitoyltransferase that decreases hepatic ceramide levels(Holland et al., 2007), would affect growth in an activated β-catenin-dependent manner. We found that 10 μM myriocin treatment modestly affected anchorage independent growth of *4x-Ala-CTNNB1* without doxycycline, but fully blocked the entire growth benefit of activated β-catenin when doxycycline was present (Fig 1H). Our cell line data suggests that ceramide synthesis contributes to the growth-promoting effects of mutant β-catenin.

### Quantification of phospholipid synthesis fluxes in human hepatocytes

The lipidomics experiments described above highlighted steady-state changes in ceramides, SM, PS, and PG lipids that could be a common manifestation of global changes in phospholipid and sphingolipid metabolism (Fig 2A). To quantify changes in phospholipid metabolism that occur upon β-catenin activation, we employed stable isotope tracer analysis. Tracer analysis provides quantitative information about the contributions of distinct pathways to the production of specific metabolites and the dynamics of each species. The use of deuterated tracers has been well-documented for quantifying phospholipid fluxes in cell culture and in healthy animals (Zhang et al., 2017). We employed a lipid head-group tracer strategy using deuterium labeled ethanolamine (4-^2^H ethanolamine), serine (3-^2^H serine), or choline (9-^2^H choline) to quantify phospholipid synthesis flux across the three major classes of *de novo* phospholipid synthesis (phosphatidylethanolamine, phosphatidylserine, and phosphatidylcholine respectively) (Fig 2A). *4x-Ala-CTNNB1* cells were cultured in standard media for 48 hours supplemented with one of the phospholipid head-group tracers. Metabolites were extracted and isotope incorporation in both aqueous and organic phase extractions was quantified by LC-MS. Lipid isotope enrichment was quantified using electrospray ionization in positive mode of parental non-fragmented species. Using a similar approach common in protein labeling experiments (Wolfe et al., 2005), we then calculated the fractional synthesis rate (FSR) or turnover of each labeled lipid species based on the observed labeling fraction normalized to the tracer exposure:

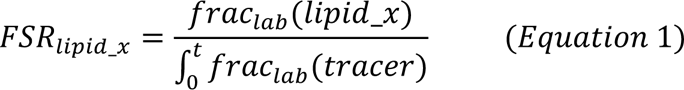

where t equals time (hrs) and *frac_lab_*(tracer) is the fractional labeling of the introduced tracer determined by media or plasma sampling over the course of the experiment.

**Figure 2:**
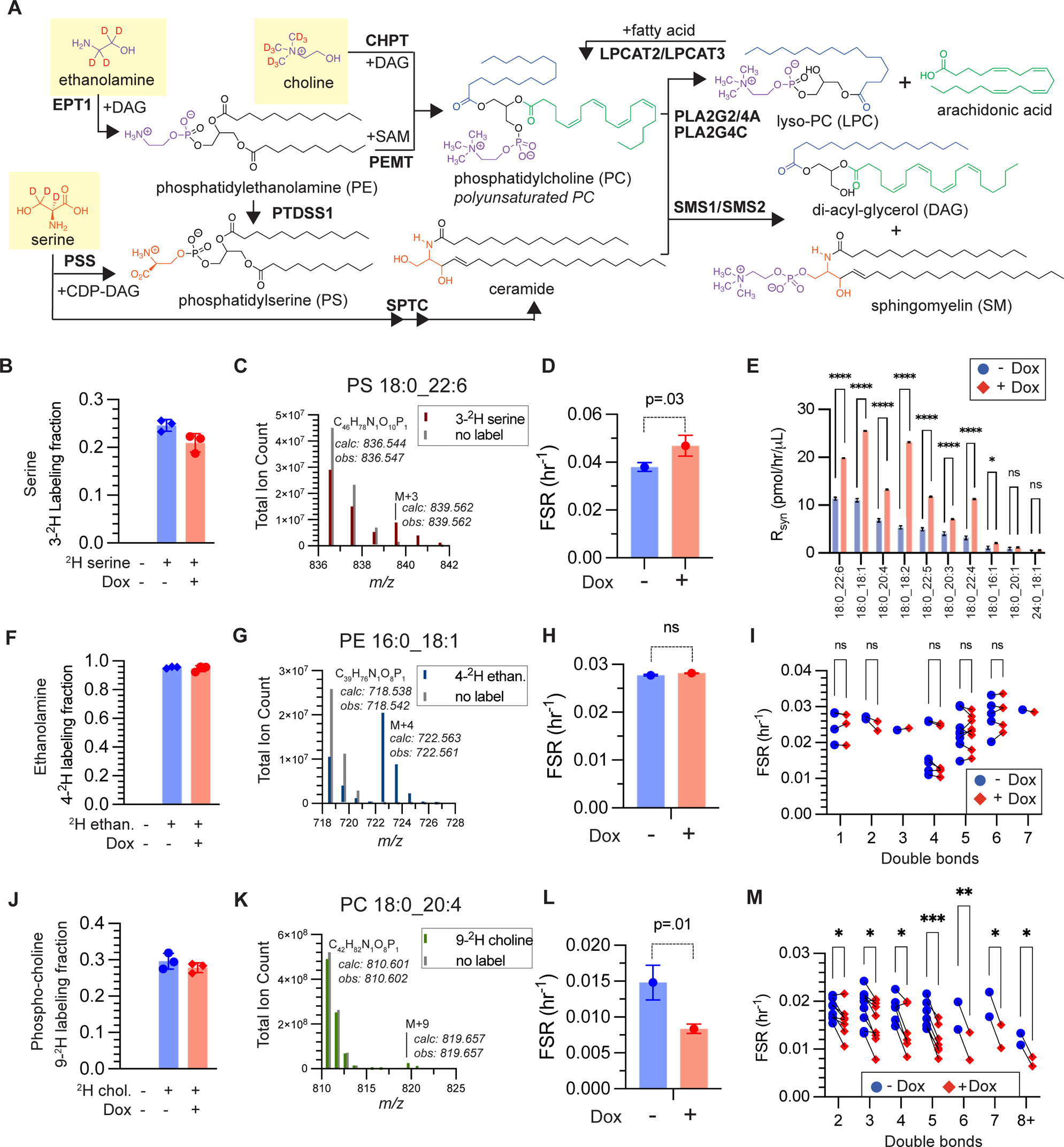
A method for quantifying phospholipid synthesis fluxes. A. A tracer strategy to label major phospholipid and sphingolipid classes. Major enzyme names are in bold. B. Recovered labeling fraction of serine from THLE-2 cells +/− Dox in *4x-Ala-CTNNB1* overexpression construct treated with 3-^2^H serine (n=3, mean±stdev). C. Representative mass spectra of PS(18:0_22:6) from cells with and without ^2^H serine label. D. Fractional synthesis rate (FSR) of PS(18:0_22:6) from THLE-2 cells +/− Dox (n=3, mean±stdev). E. Absolute synthesis rates (R_syn_) of PS lipids from THLE-2 cells +/− Dox (mean±stdev). F. Recovered labeling fraction of ethanolamine from THLE-2 cells +/− Dox treated with 4-^2^H ethanolamine (n=3, mean±stdev). G. Representative mass spectra of PE(16:0_18:1) from cells with and without ^2^H ethanolamine label. H. FSR of PE(16:0_18:1) from THLE-2 cells +/− Dox (n=3, mean±stdev). I. FSR of PE lipids from THLE-2 cells +/− Dox by total fatty acid unsaturation (n=3). J. Recovered labeling fraction of phosphocholine from THLE-2 cells +/− Dox treated with 9-^2^H choline (n=3, mean±stdev). K. Representative mass spectra of PC(18:0_20:4) from cells with and without ^2^H choline label. L. FSR of PC(18:0_20:4) from THLE-2 cells +/− Dox (n=3, mean±stdev). M. FSR of PC lipids from THLE-2 cells +/− Dox by total fatty acid unsaturation (n=3). * p<.05, ** p<.01, *** p<.001, **** p<.0001. For D, H, and L, Student’s unpaired t-test. For E, multiple unpaired t-tests; for I and M, multiple paired t-tests; corrected for multiple comparisons using the Holm-Šídák method

Enrichment of 3-^2^H serine was not significantly different between *4x-Ala-CTNNB1* cells with and without doxycycline (Fig 2B), indicating that the uptake and metabolism of the tracer itself was not different between experimental and control groups. 3-^2^H serine labeling of representative lipid PS(18:0_22:6) was clearly evident above background, and the FSR was increased with activated β-catenin expression (Fig 2C). Across nearly all quantified PS species, FSR was increased upon doxycycline treatment (Supplemental Fig 2A). Combining FSR and absolute abundance data, we determined the synthesis rate (R_syn_) of each PS lipid. For 8 of the 10 quantified PS lipids, R_syn_ was significantly increased by doxycycline treatment (Fig 2E) indicating enhanced flux through the phosphatidylserine synthetase reaction. In contrast, we did not observe differences in serine incorporation into ceramides, indicating that the observed increase in ceramides came from changes to degradation pathways, and not de novo sphingolipid synthesis (Supplemental Fig 2B).

As with 3-^2^Hserine, 4-^2^H ethanolamine labeling of cells was unaffected by doxycycline and observable in lipids species like PE(16:0_18:1) (Fig 2F,G). However, labeling of PE lipids and the calculated turnover rate was not different between *4x-Ala-CTNNB1* cells treated with doxycycline compared to those without (Fig 2H, Supplemental Fig 2C). We did not observe any changes in FSR upon β-catenin activation when PE lipids were sorted by saturation (Fig 2I). Under the growth conditions we used for THLE-2 cells, we did not observe conversion of PE lipids into PC lipids.

Cellular labeling with 9-^2^H choline resulted in a strong enrichment of 9-^2^H phosphocholine (Fig 2J). Labeling of phosphatidylcholine lipid species PC(18:0_20:4) was evident in the M+9 mass (Fig 2K). Unlike with PE lipids, for most PC lipid species, FSR was reduced upon doxycycline treatment and β-catenin activation (Fig 2L, Supplemental Fig 2D). The reduction in PC *de novo* synthesis flux was observed across all levels of lipid saturation (Fig 2M). Total synthesis (R_syn_) of PC lipids was modestly but significantly reduced in *4x-Ala-CTNNB1* cells (Supplemental Fig 2E).

Together, our tracing data indicate that activated β-catenin in hepatocytes promotes significant changes in phospholipid metabolism exemplified by the enhancement of PS synthesis flux while simultaneously inhibiting PC synthesis flux through the CDP-choline pathway.

### Activated β-catenin leads to lipidome dysregulation in a transgenic zebrafish model of HCC

Metabolism is highly context dependent, and standard cell culture conditions represents only one possible set of myriad in vivo conditions. To understand how activated β-catenin impacts lipid metabolism in whole organisms, we turned to a zebrafish HCC model. Transgenic zebrafish expressing hepatocyte-specific activated β-catenin (Tg-ABC) from *Xenopus laevis* (S33A, S37A, T41A, and S45A) develop HCC that morphologically and genetically recapitulates key features of human HCC (Evason et al., 2015). Unlike mouse HCC models driven by activated β-catenin, Tg-ABC zebrafish do not require co-expression of a second oncogene or treatment with carcinogens to induce tumors (Harada et al., 2004; Qiao et al., 2018; Tao et al., 2017).

We performed LC-MS lipidomic analysis and GC-MS polar metabolite analysis on male and female Tg-ABC HCC livers alongside non-tumor livers from non-transgenic, sex-matched, control siblings (non-Tg). Principal component analysis (PCA) of polar metabolites did not partition livers by either sex or tumor status (Supplemental Fig 3A). Univariate analysis showed that individual polar metabolites, including free fatty acids such as palmitic acid and steric acid as well as glycolytic intermediate 3-phosphoglycerate, increased in Tg-ABC livers compared to non-Tg livers (Supplemental Figure 3B). Pyruvate was significantly downregulated as was glycerol-3-phosphate. While the overall polar metabolome of Tg-ABC livers was not significantly distinct from the non-Tg animals, elevated free fatty acids and reduced pyruvate suggest a reduction in glycolytic flux and potentially an increase in lipolysis for fatty acid oxidation.

To determine if glycolytic flux was in fact reduced in activated β-catenin-driven tumors, we injected zebrafish interperitoneally with uniformly labeled (U)-^13^C glucose and collected livers after 2 hours. The total 6-^13^C-labeled glucose fraction was not different between non-Tg and Tg-ABC livers indicating no significant change in glucose uptake (Supplemental Fig 3C). In contrast, the 3-^13^C-labeled lactate fraction was reduced by greater than 50% (adjusted p value 0.02), showing a highly reduced rate of glycolysis (Supplemental Fig 3C). The contribution of glucose and glutamine to TCA metabolism was not different between Tg-ABC and non-Tg livers, and collectively they only accounted for ∼25% of TCA substrate utilization (Supplemental Fig 3D). Overall, our polar tracing and metabolomics data showed notable differences in glycolytic metabolism in Tg-ABC livers compared to non-Tg, but few other significant alterations.

In contrast, metabolite differences between non-Tg and Tg-ABC livers were clearly apparent when we analyzed the lipidome. PCA of total LC-MS quantified lipid abundances partitioned Tg-ABC HCC and non-Tg liver lipidomic profiles and additionally showed clear sex-specific separation (Fig 3A). The principal component associated with sex (PC1) accounted for 3 times the variance of PC2, which more closely matched tumor status. Henceforth, we analyzed lipidomic data from Tg-ABC and non-Tg zebrafish in a sex-specific manner.

**Figure 3:**
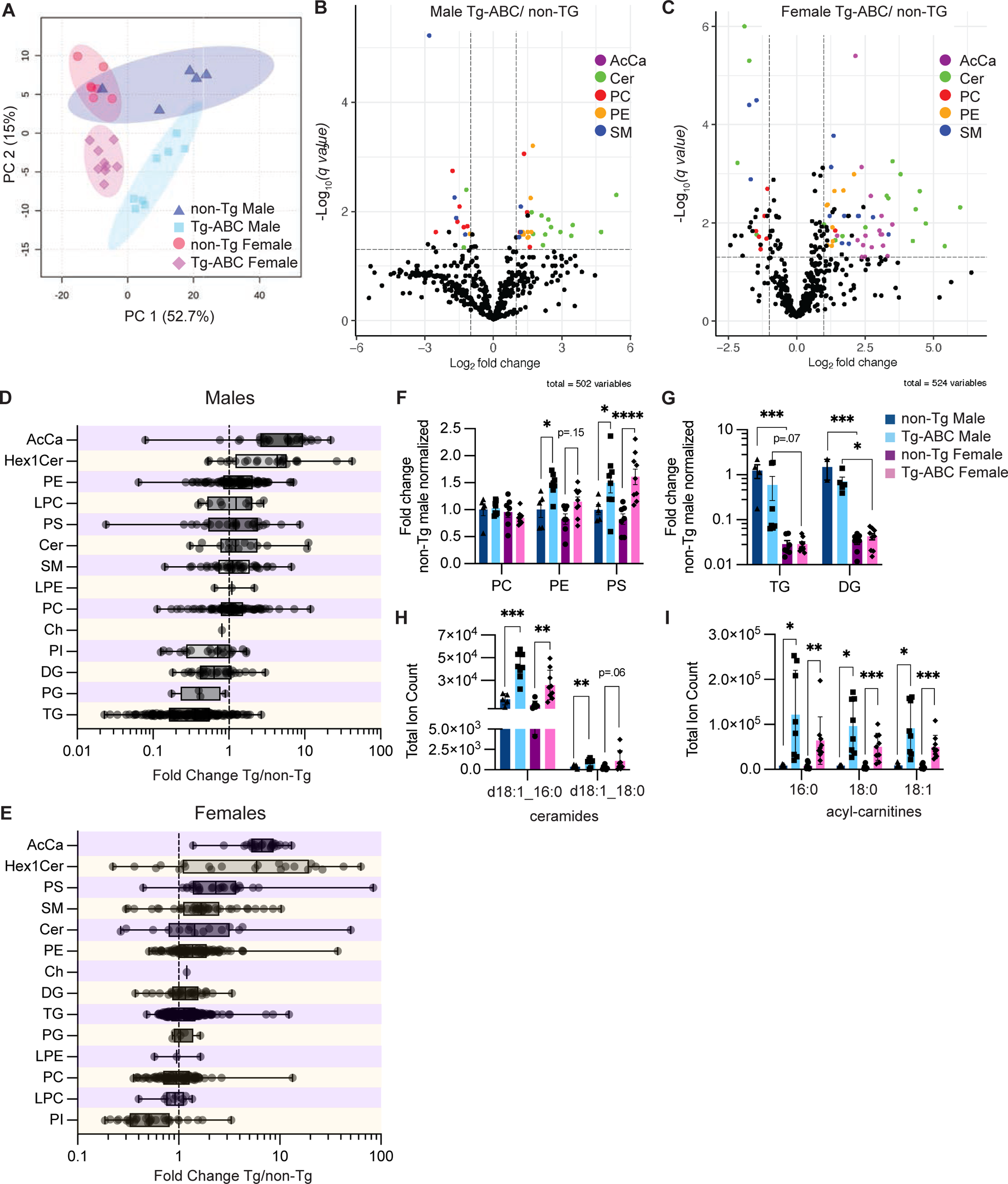
Tg-ABC zebrafish livers show sex-specific remodeling of lipid metabolism. A. Principal component analysis of all quantified lipid species extracted from livers of male and female, non-Tg and Tg-ABC zebrafish (n=5-9). B,C. Volcano plots showing change in abundance for individual lipid species from livers of non-Tg and Tg-ABC, male (B) and female (C) zebrafish (n=5-9). Y axis is the −Log_10_q using a multiple-comparisons corrected false-discovery rate of 5%. D,E. Fold changes in lipid species by class. Lipid classes are defined in Fig 1. Fold change is average change between livers from Tg-ABC (Tg) and non-Tg fish (n=5-9). F. Change in total abundance of phospholipid species from non-Tg and Tg-ABC, male and female zebrafish livers normalized to non-Tg male abundance (n=5-9, mean±SEM). G. Change in total abundance of triglycerides (Tg) and diglycerides (Dg) by sex and genotype (n=3-9, mean±SEM). H. Total ion counts of select d18:1 ceramides species by sex and genotype (n=5-9, mean±SEM). I. Total ion counts of select acyl-carnitine species by sex and genotype (n=5-9, mean±SEM). For F-I, * p<.05, ** p<.01, *** p<.001, **** p<.0001 by 2-way unpaired ANOVA, corrected for multiple comparisons by Tukey’s method.

Male Tg-ABC zebrafish livers showed significant dysregulation of 46 lipids out of 502 quantified (Fig 3B). Similar to our observations in *4x-Ala-CTNNB1* cells, ceramides and hexosylceramides (Hex1Cer) were elevated in male Tg-ABC livers compared to non-Tg control siblings. PE and PC lipids including PC(18:0_20:4) and PE(18:0_20:4) were also present among the most up- and down-regulated lipids (20/46 were PE or PC species). The lipid class with the greatest percentage of elevated members was acylcarnitines. Unlike in the *4x-Ala-CTNNB1* cells, SM lipids were not significantly downregulated and individual species were among the most upregulated lipid metabolites (Fig 3B,D). Storage lipids accounted for the most downregulated lipid classes in male Tg-ABC livers, including triglycerides followed by PG and DG lipids (Fig 3D).

Lipidome remodeling was even more significant in female Tg-ABC zebrafish livers compared to non-Tg livers with 86 lipids out of 524 quantified significantly altered (Fig 3C). As in *4x-Ala-CTNNB1* cells and male Tg-ABC zebrafish, ceramides were clearly upregulated by activated β-catenin expression. Acyl-carnitine species, PE lipids, and SM lipids were significantly increased in abundance. At the class level, the increase in PS lipids was greater than for other phospholipids (Fig 3E). Unlike in male fish, storage lipids (TG, GD and PG) were not significantly altered, and the only downregulated class was PI lipids (Fig 3E).

We directly compared changes in major lipid classes between male and female zebrafish. For storage lipids, there was a clear sexual dimorphism: non-Tg males had ∼35 fold higher concentrations of TG and DG lipids compared to non-Tg females (Fig 3G). These sex-dependent differences in TG and DG levels were attenuated but not eliminated in Tg-ABC livers.

Both sexes showed similar changes in phospholipid abundances between non-Tg and Tg-ABC livers: Tg-ABC livers showed no differences in PC lipids as a class, but had increased levels of PE and PS lipids (Fig 3F). Male and female zebrafish also demonstrated conserved changes in specific key acylcarnitine and ceramide species. The bioactive ceramides Cer(d18:1_16:0), Cer(d18:1_18:0), Cer(d18:1_18:1), were all significantly increased in male and female Tg-ABC livers (Fig 3H). Similarly, acylcarnitine 16:0, 18:0, and 18:1 were all increased in Tg-ABC compared to non-Tg livers (Fig 3H). To explore how these consistent alterations in the tumor lipidome may arise in livers with sexually dimorphic patterns of lipid metabolism, we analyzed RNA expression data and performed lipid tracing analysis.

### Consistent alterations in zebrafish and human HCC lipid metabolic gene networks

Inferences about metabolism can be made from the analysis of expression levels of genes encoding key metabolic enzymes. To understand the sources of HCC induced lipid metabolic changes, we re-analyzed publicly available gene expression data from male (Kalasekar et al., 2019) and female (Evason et al., 2015) Tg-ABC HCC zebrafish and non-transgenic, sex-matched control siblings, as well as human HCC data from The Cancer Genome Atlas Liver Hepatocellular Carcinoma (TCGA-LIHC)(Cancer Genome Atlas Research Network, 2017). In addition to analyzing paired HCC and non-tumor samples from all human HCC regardless of mutational profile (TCGA-All), we also performed a nested analysis of TCGA-LIHC data to compare tumors with mutated *CTNNB1* to those without (TCGA-ABC). We validated our analysis by identifying GO biological processes related to Wnt signaling as being significantly enriched in male and female zebrafish β-catenin-driven HCC (Tg-ABC) and in human *CTNNB1-*mutated HCC (TCGA-ABC), but not in the broader human HCC analysis (TCGA-All) (Table 1). We found that “lipid metabolic process” (GO:0006629) and “cellular lipid metabolic process” (GO:0044255) were significantly enriched in male and female Tg-ABC HCC zebrafish, as well as in human patients with HCC (TCGA-All) (Table 1). These were the only metabolism associated GO terms significantly enriched across all three HCC datasets. Comparing patients with mutated *CTNNB1* to those without (TCGA-ABC), “lipid metabolic process” (GO:0006629) and “cellular lipid metabolic process” (GO:0044255) were further significantly enriched (Table 1). We performed targeted transcriptomic analysis of key enzymes in phospholipid metabolism and identified substantial similarities in dysregulated genes between male Tg-ABC HCC zebrafish, female Tg-ABC HCC zebrafish, TCGA-All, and TCGA-ABC (Fig 4). Three of the four datasets showed that HCC was associated with significant upregulation of at least one isoform of *carnitine palmitoyltransferase* (*CPT)*, which encodes a key enzymatic regulator of β-oxidation. In all datasets, there was a significant downregulation in *phosphatidylethanolamine N-methyltransferase* (*PEMT)*, which encodes the enzyme catalyzing the conversion of PE to PC, and significant dysregulation in at least one isoform of *sphingomyelin phosphodiesterase* (*SMPD)*, which encodes the enzyme catalyzing the conversion of sphingomyelin to ceramide and PC (Fig 4, Sup Table 1). Together, these results highlight substantial lipid metabolic changes in activated β-catenin-driven HCC in zebrafish and in human patients, especially with respect to fatty acid oxidation, phospholipid and sphingolipid metabolism.

**Figure 4:**
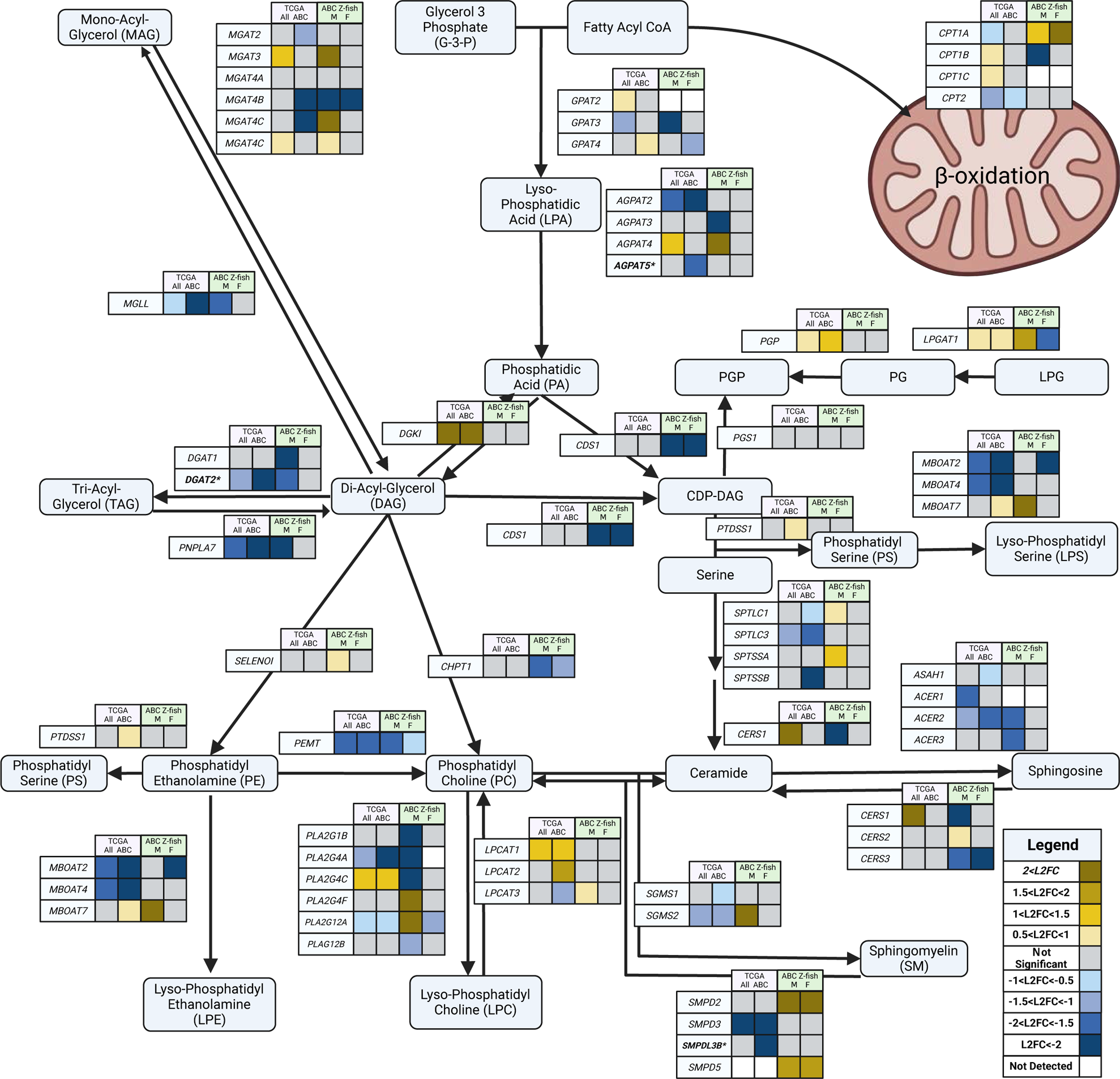
Transcriptomics Reveals Dysregulated Lipid Metabolic Pathways Between Human and Zebrafish ABC-driven HCC. Transcriptomics was analyzed using DESeq2 (v1.34.0) with GRCh38 and GRCz11 for Human and Zebrafish respectively with padj<0.05 and an absolute L2FC >0.5. Genes that were significantly dysregulated between the tumor of patients with mutated *CTNNB1* to tumor of patients with non-mutated *CTNNB1* are bolded with an asterisk. PGP: Phosphatidylglycerol phosphate; PG: Phosphatidylglycerol; LPG: Lysophosphatidylglycerol.

**Table 1:**
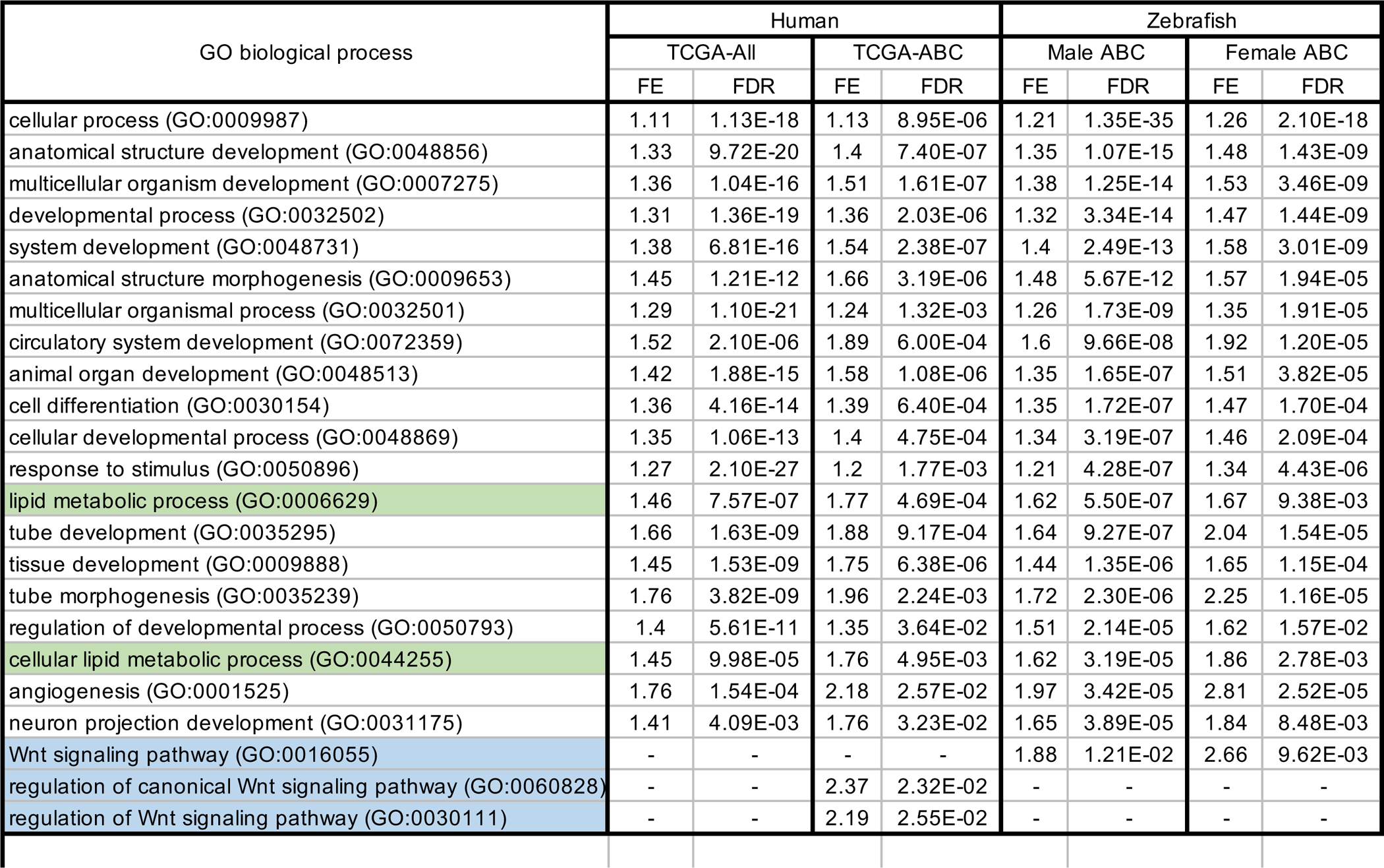
GeneOntology Analysis Reveals Dysregulated Lipid Metabolism Between Human and Zebrafish ABC-driven HCC. Transcriptomics was analyzed using DESeq2 (v1.34.0) with GRCh38 and GRCz11 for Human and Zebrafish respectively with padj<0.05 and an absolute L2FC >1.0. Human analyses were performed for TCGA-ALL with Tumor vs Non-Tumor for all paired samples of TCGA-LIHC. TCGA-ABC was Tumor vs Non-Tumor specifically for 11 male patients with documented mutations in *CTNNB1*. Zebrafish analyses were performed by sequencing tissue from adult zebrafish liver from transgenic zebrafish (Tg-ABC) compared to non-transgenic, sex-matched control siblings. Table was collated with Identical GO terms and sorted by male transgenic Tg-ABC zebrafish.

### Quantification of phospholipid fluxes reveals female-specific suppression of *de novo* PC synthesis pathway in tumors

Based on the centrality of phospholipid metabolism to many of the observed changes in the steady-state lipidome, we adopted our phospholipid isotope tracing strategy from cells to live zebrafish to quantify lipid fluxes and identify key changes in reaction fluxes (Fig 2). We implemented our headgroup tracer strategy for use in zebrafish by performing a single intraperitoneal injection (IP) with deuterium-labeled 9-^2^H choline or 4-^2^H ethanolamine and then extracted lipids from livers 24 hours later (Fig 5A). We were unable to observe lipid incorporation of 3-2H serine by this single-injection method to introduce isotope. Uptake and incorporation of tracers was detected by recovering labeled phospholipids in both Tg-ABC HCC zebrafish and non-Tg control siblings, and isotope labeling from MS spectra was calculated for individual phospholipid species.

**Figure 5:**
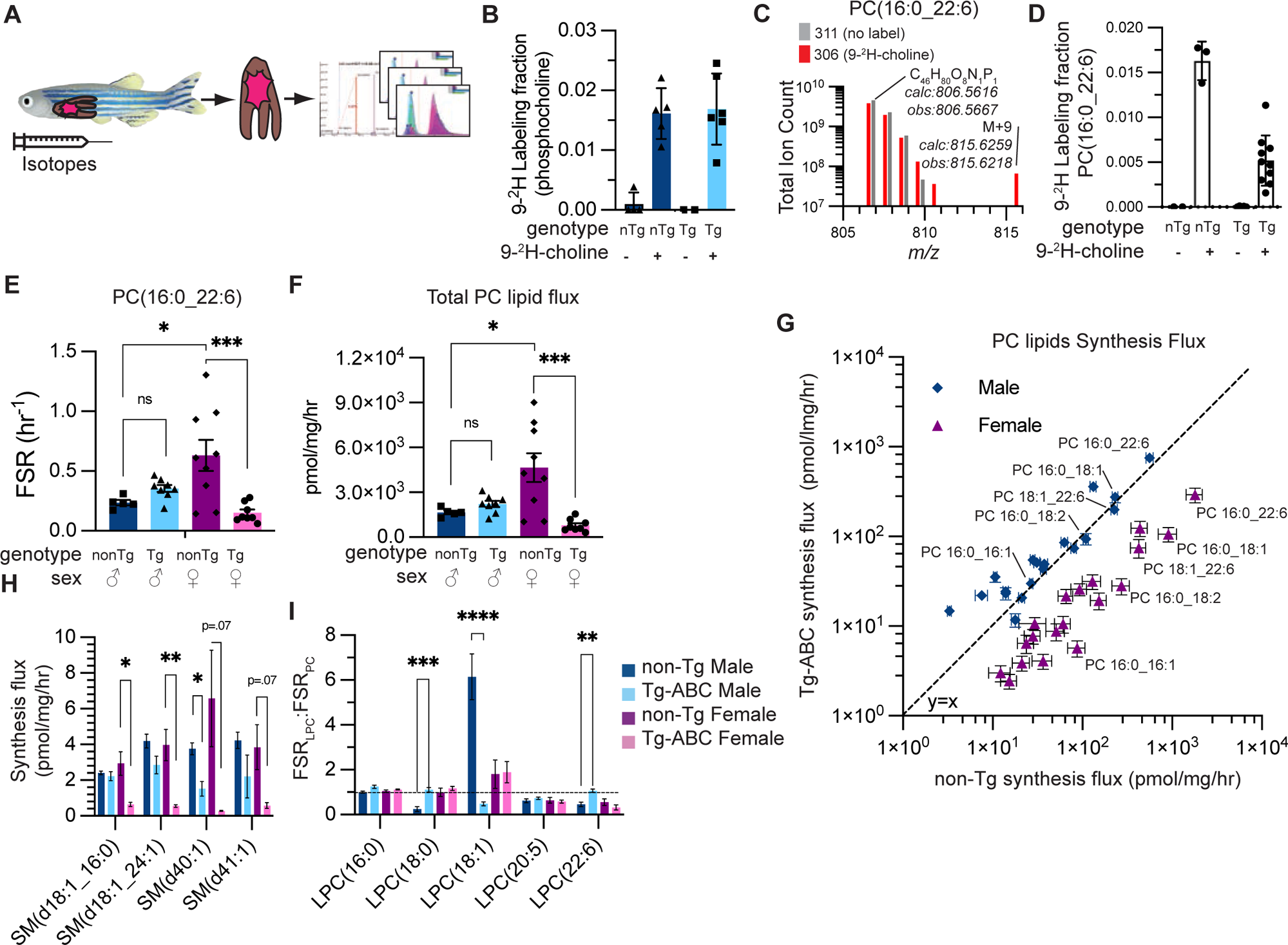
A method for quantifying phospholipid fluxes *in vivo* identifies a female-specific suppression of phosphatidylcholine synthesis in liver tumors. A. An overview of the strategy to introduce stable isotope precursors into zebrafish to label lipids. B. Recovered labeling fraction of phosphocholine in non-Tg (nTg) and Tg-ABC (Tg) zebrafish livers with or without 9-^2^H choline injection (n=2-6, mean±SEM). C. MS spectra of PC(16:0_22:6) in zebrafish livers from animals injected with PBS or 9-^2^H choline 24 hrs earlier. D. 9-^2^H labeling fraction of PC(16:0_22:6) in nTg and Tg zebrafish livers with or without 9-^2^H choline injection (n=2-10, mean±SEM). E. FSR of PC(16:0_22:6) as calculated from 9-^2^H choline incorporation in non-Tg and Tg, male and female zebrafish livers (n=5-9, mean±SEM). F. Total quantified synthesis rate of all PC lipids in zebrafish livers by sex and genotype (n=5-9, mean±SEM). G. Absolute synthesis rates of select PC lipids for non-Tg (x-axis) and Tg-ABC (y-axis) zebrafish livers by sex (n=5-9, mean±SEM). Select individual species are named. H. Absolute synthesis rate of sphingomyelin lipids (SM) as calculated from 9-^2^H choline incorporation and absolute abundance by sex and genotype (mean±SEM). I. FSR of LPC lipids normalized to FSR of PC lipid precursor by sex and genotype (n=5-9, mean±SEM). For E, F, H, and I, * p<.05, ** p<.01, *** p<.001, **** p<.0001 by 2-way unpaired ANOVA, corrected for multiple comparisons by Tukey’s method.

To monitor 9-^2^H choline enrichment in livers, we again relied on the first committed choline metabolite, 9-^2^H phosphocholine. We observed 9-^2^H phosphocholine enrichment 24 hours after injection in non-Tg and Tg-ABC livers injected with 9-^2^H choline compared to saline controls (Fig 5B). To understand total tracer exposure in these animal experiments, we monitored enrichment of 9-^2^H phosphocholine extracted from livers at multiple timepoints after injection. Tracer exposure fit first order kinetics over the 24 hours of the experiment (Supplemental Fig S5A), consistent with a model of tracer exposure based on the assumption of a non-saturating bolus of tracer. In Tg-ABC zebrafish compared to non-Tg sibling controls, we observed an increased rate of decay with slightly higher initial enrichment values, leading to a modestly higher total exposure. All subsequent calculations of lipid turnover fluxes were normalized to these genotype-dependent differences in tracer exposure.

The identification and quantification of isotope-labeled liver phospholipids was performed as done in the cell culture experiments. Total observed isotopic enrichment of 9-^2^H labeled phospholipids was lower than that from cell culture samples and in the range of 0.5 to 1.5%, but clearly separated from background natural isotopic abundance (Fig 5C,D). Among female zebrafish, representative phospholipid PC(16:0_22:6) was less labeled in Tg-ABC compared to non-Tg control siblings (Fig 5D). We observed isotopic labeling from 9-^2^H choline above background in a set of 27 abundant identified PC lipid species in both Tg-ABC and non-Tg zebrafish livers.

There were significant sex- and tumor-specific differences in PC synthesis rates (Fig 5E, Supplemental Fig S5B). Non-Tg female zebrafish had ∼2-3 fold greater rates of *de novo* PC lipid synthesis compared to non-Tg male zebrafish across all quantified species, including representative PC(16:0_22:6) (Fig 5E). PC synthesis flux from choline was unaltered or modestly increased in Tg-ABC livers in male zebrafish. In female fish, this pathway was significantly downregulated in Tg-ABC livers compared to non-Tg control siblings. Overall, female Tg-ABC zebrafish had levels of PC synthesis flux that were lower than their male Tg-ABC counterparts (Supplemental Fig 5B).

We calculated the absolute synthesis rate (R_syn_) of each PC lipid species. The total sum of PC synthesis flux through the CDP-choline pathway was significantly reduced in female Tg-ABC but not male Tg-ABC zebrafish (Fig 5F). In total, we were able to quantify absolute synthesis rates for 19 PC lipids (Fig 5G). As expected from the modest changes in FSR, total *de novo* PC synthesis flux in male Tg-ABC fish was nearly identical to that in non-Tg fish with significant deviations only in very low-abundance species. In contrast, for females, Tg-ABC flux was significantly repressed across PC lipids represented over 3 orders of magnitude in abundance.

We next identified 9-^2^H choline incorporation into SM lipids, which are produced from PC lipids by the activity of sphingomyelin synthases (Fig 2A). We used the 9-^2^H enrichment fraction in 4 SM species to calculate synthesis rate from *de novo* choline incorporation (Fig 5H). As SM lipid tails do not originate from PC lipids, we were unable to normalize the observed labeling by the direct precursor labeling. As such, the synthesis flux we report here reflects the combined activities of PC and SM synthesis reactions. Among quantifiable SM species, we did not observe a significant difference in SM synthesis rates between female and male non-Tg zebrafish or between male Tg-ABC and non-Tg zebrafish (Fig 5H). In contrast, we observed reduced SM synthesis in female Tg-ABC zebrafish compared to non-Tg control siblings.

Finally, we quantified the production of LPC lipids from PC lipids, which are produced from PC lipids by the activity of phospholipases (Fig 2A). As each LPC lipid contains a lipid tail that identifies a set of related potential precursor PC lipids, we determined the FSR for each LPC lipid and compared that to the average FSR of the potential PC precursors to give a FSR_LPC_:FSR_PC_ ratio (Fig 5I, Supplemental Fig 5C). An FSR_LPC_:FSR_PC_ ratio equal to 1 indicates that the observed phospholipase reaction to generate the LPC species matches the production of the precursor PC lipid pool; an FSR_LPC_:FSR_PC_ ratio greater than 1 implies that the LPC species is being generated faster than the PC precursor; and an FSR_LPC_:FSR_PC_ ratio less than 1 indicates slower LPC lipid synthesis rates. For most conditions tested, normalized FSR_LPC_:FSR_PC_ ratios were ∼1 (Fig 5I). Among female fish, there were no differences in FSR_LPC_:FSR_PC_ ratios between Tg-ABC and non-Tg livers. Among male fish, however, there were significant differences in FSR_LPC_:FSR_PC_ ratios for three LPC lipid species (18:0, 18:1, and 22:6). Non-Tg male zebrafish had a high rate of production of the LPC species 18:1, which was reduced nearly 10-fold in Tg-ABC zebrafish. Non-Tg male zebrafish had a low rate of production of the LPC species 18:0 and 22:6, which was significantly increased in Tg-ABC zebrafish.

In summary, we found that non-Tg female zebrafish showed significantly greater *de novo* PC lipid synthesis from choline than non-Tg male zebrafish. During β-catenin-driven liver tumorigenesis, female zebrafish demonstrated significantly decreased *de novo* PC lipid synthesis from choline while males had no significant change in this pathway. These tracing data also highlight significant sex-specific differences in phospholipase activity in both Tg-ABC and non-Tg zebrafish.

### Male Tg-ABC fish are deficient in lipid methylation

To complete our analysis of PC lipid metabolism, we next utilized 4-^2^H ethanolamine tracer to examine PC lipid synthesis through PE methylation, the other pathway of *de novo* synthesis. We again performed a single intraperitoneal injection of fish, waited 24 hours, and then excised livers for LC-MS analysis. After 24 hours, total 4-^2^H enrichment in ethanolamine was between 5-10%, and higher in non-Tg male zebrafish than in non-Tg female zebrafish (Fig 6A). We used additional fish to determine the approximate kinetics of ethanolamine metabolism and observed much slower decay compared to choline tracer (Supplemental Fig 6A). Using that data, we determined a total exposure for each fish as we had done with choline. 4-^2^H labeled phosphatidylethanolamine (PE) was readily detected and present at a higher enrichment fraction than 9-^2^H PC, reflecting both the increased tracer exposure and smaller pool size of PE lipids compared to PC lipids (Fig 6B). We computed synthesis rates by multiplying the exposure normalized labeling fraction (FSR) by the absolute abundance of the lipid species (Supplemental Fig 6B, Fig 6C). For a set of abundant identified PE lipids, a common pattern was present in most species: Total PE lipid synthesis rates were similar between non-Tg male and female zebrafish but elevated in Tg-ABC zebrafish. This pattern was exemplified by synthesis rates of PE 16:0_20:5 (Fig 6C). This labeling pattern indicates enhanced *de novo* synthesis of PE lipids from ethanolamine catalyzed by ethanolaminephosphotransferase (*SELENOI*). This is consistent with the observed steady-state increases in total PE lipids in Tg-ABC fish of both sexes and the increased gene expression of *SELENOI* in male fish (Fig 4).

**Figure 6:**
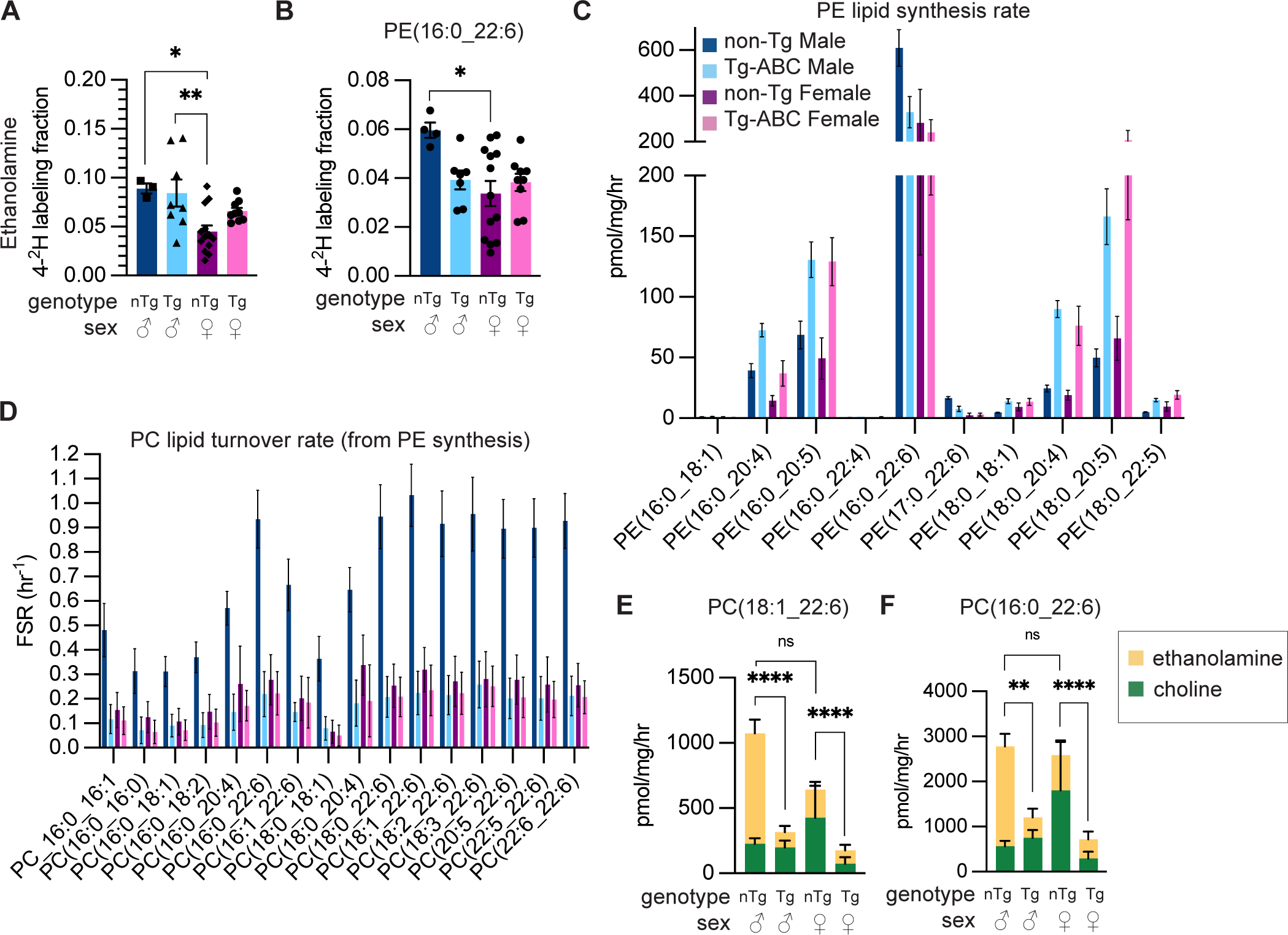
Male zebrafish suppress PE lipid methylation to limit PC lipid synthesis in β-catenin-driven liver tumors. A. Liver labeling fraction of ethanolamine in non-Tg (nTg) and Tg-ABC (Tg), male and female zebrafish livers 24 hrs after 4-^2^H ethanolamine injection (n=4-13, mean±SEM). B. 4-^2^H labeling fraction of PE(16:0_22:6) in zebrafish livers by genotype and sex (n=4-13, mean±SEM). C. Absolute synthesis rates for PE lipids as calculated from 4-^2^H ethanolamine incorporation in zebrafish livers by genotype and sex (mean±SEM). D. FSR of PC lipids as calculated from 4-^2^H ethanolamine incorporation in zebrafish livers by genotype and sex (n=5-9, mean±SEM). E, F. Total PC synthesis flux for representative PC lipids as determined by ethanolamine and choline tracers by sex and genotype. Significance is based on the total synthesis flux from both quantified pathways (mean±SEM). For A, B, E, and F, * p<.05, ** p<.01, *** p<.001, **** p<.0001 by 2-way unpaired ANOVA, corrected for multiple comparisons by Tukey’s method.

PE lipids are the source of a significant fraction of PC lipids via the methylation activity of PEMT to convert PE into PC. We observed 4-^2^H enrichment in PC lipids from zebrafish livers injected with 4-^2^H ethanolamine (Fig 6D). The calculated FSR data revealed a significantly elevated rate of PC production via PEMT in non-Tg male zebrafish compared to non-Tg female or Tg-ABC male animals, a pattern not present in the analysis of ethanolamine incorporation into PE lipids. From these data, we calculated an absolute synthesis flux of PC lipids through the PEMT pathway and combined that with our CHPT1 synthesis flux calculation (Supplemental Figure 6C,D). Examining these analyses concurrently, we observed that total PC synthesis flux was roughly equal between non-Tg male and female livers but was significantly reduced in Tg-ABC livers in both sexes (Fig 6E,F and Supplemental Fig 6D). Thus, both male and female Tg-ABC zebrafish showed decreased PC flux, though this global metabolic change was mediated by alterations in different pathways in the two sexes: The CDP-choline pathway was inhibited in female Tg-ABC zebrafish while the PE methylation pathway was inhibited in male Tg-ABC zebrafish (Supplemental Figure 6E).

## Discussion

Changes in lipid metabolism are widespread in tumors (Broadfield et al., 2021). Identifying and linking changes in metabolic activity with specific oncogenic signaling events creates an opportunity to target tumor lipid metabolism therapeutically. In this work, we present a detailed description of steady-state changes and dynamic phospholipid flux alterations induced by activated β-catenin from two independent model systems of HCC. Changes in common to the two systems included a steady-state increase in ceramides and acylcarnitines, a decrease in triglycerides, and significant remodeling of the phospholipidome. Our tracing data reveals a global suppression of *de novo* phosphatidylcholine biosynthesis achieved by sex-dependent mechanisms in adult zebrafish.

In this study, we utilized a zebrafish model of HCC driven by a *CTNNB1* transgene in order to study tumor lipid metabolism (Evason et al., 2015). While zebrafish have historically been utilized for their strengths in developmental studies and chemical screens, adult zebrafish have gained popularity as a model to study metabolic processes in both normal physiology and cancer, facilitated by new experimental approaches including dietary intervention (Chen et al., 2018; Forn-Cuní et al., 2015; Landgraf et al., 2017; Oka et al., 2010), intraperitoneal injections (Kinkel et al., 2010), and tissue isolation for mass-spectrometry (Bosworth et al., 2005; Choi et al., 2015; Purushothaman et al., 2019; Shrader et al., 2003; Tay et al., 2006). Naser et al. recently and elegantly quantified metabolic fluxes in a zebrafish model of melanoma using ^13^C glucose immersion to introduce isotope into the system (Naser et al., 2021). Here we extend this method to intraperitoneal injection and a broader range of isotopically labeled nutrients in an HCC model, demonstrating that is feasible to obtain stable incorporation of ^2^H into lipids species in both healthy and diseased zebrafish liver.

When considering the effects of activated β-catenin on the steady-state lipidome, we observed the most striking changes in metabolites and gene expression related to fatty acid β-oxidation: increased acylcarnitines in *4x-Ala-CTNNB1* THLE-2 cells and male and female Tg-ABC zebrafish, as well as decreased triglycerides and diglycerides in *4x-Ala-CTNNB1* THLE-2 cells and male Tg-ABC zebrafish (Fig 1 and 3). These changes in acylcarnitines and storage lipids were accompanied by increased *CPT* expression in dox-treated *4x-Ala-CTNNB1* THLE-2 cells, Tg-ABC zebrafish and human HCC. Our observations are consistent with prior reports that activation of β-catenin signaling via Wnt ligand activation or APC loss drives fatty acid oxidation in osteoblasts (Frey et al., 2018) and mouse HCC (Senni et al., 2019). We quantified glucose metabolism in zebrafish HCC and observed a reduction in glycolytic flux. These data are consistent with reported ^18^FDG-PET data from mouse (Chafey et al., 2009) and human (Gougelet et al., 2019) demonstrating that activated β-catenin tumors showing poor glucose uptake. Altogether these results support a model in which activated β-catenin drives a shift away from glucose metabolism and increases reliance on fatty acid oxidation for growth, highlighting the therapeutic potential of blocking fatty acid oxidation in mutant *CTNNB1* HCC.

We found significantly increased levels of ceramides as a class, especially increased Cer(d18:1_16:0), in *4x-Ala-CTNNB1* THLE-2 cells and in male and female Tg-ABC zebrafish (Fig 1 and 3). Ceramides have traditionally been thought to be anti-tumorigenic by driving apoptosis in HCC (Jiang et al., 2017; Krautbauer et al., 2016), and there have been divergent reports about ceramide levels in HCC from human studies. In cohorts with large numbers of HCV (Ismail et al., 2020) and in those with hepatitis infection excluded (Krautbauer et al., 2016), liver ceramides have been reported as downregulated in HCC, whereas in a cohort with a large number of HBV infections, no changes in ceramides were reported (Lu et al., 2018). In contrast, Grammatikos et al. identified elevated serum ceramides as prognostic for differentiating HCC cases from cirrhosis in a mixed cohort of HBV- and HCV-positive and -negative patients (Grammatikos et al., 2016). Ceramides also promote progression of metabolic dysfunction-associated steatohepatitis (MASH), a well-established risk factor for HCC (Chaurasia et al., 2019). We have previously proposed that modest increases of ceramides, at levels that are insufficient to induce apoptosis, reprogram metabolism in cells in a manner that might be protumorigenic(Li et al., 2023). Two of our findings in this study provide additional evidence supporting a tumor-promoting role for ceramides in HCC. First, activated β-catenin increased bioactive ceramides in cells and in zebrafish. Second, inhibition of ceramide production via myriocin treatment decreased β-catenin-driven anchorage-independent growth. It remains possible that the β-catenin-driven increase in ceramides that we observed could be unrelated to the oncogenic effects of β-catenin or could represent a parallel compensatory response to oncogene activation.

The relationship between ceramides and Wnt/β-catenin signaling is also unclear based on published work. Two studies in cancer cells showed a consistent effect of exogenous ceramides on activating Wnt signaling in part through dephosphorylating β-catenin and increasing translocation to the plasma membrane (Azbazdar et al., 2022; Marchesini et al., 2007). Azbazdar et al. further showed that migration of SNU475 cells in a larval zebrafish assay was inhibited by pre-treatment with myriocin. On the other hand, Wnt/β-catenin signaling can also affect ceramide levels. Modulation of Wnt signaling by either addition of Wnt ligand or addition of siRNA against β-catenin has been reported to decrease or increase levels of ceramides respectively in cells or non-cancerous mice (Azbazdar et al., 2022; Popov et al., 2016). Our finding that activated β-catenin increased bioactive ceramides in both cells and in zebrafish provides support for the hypothesis that Wnt/β-catenin signaling modulates ceramide levels directly.

In both zebrafish HCC and human cell lines, we identified significant β-catenin-driven changes in steady-state phospholipid and sphingolipid abundances at both the level of individual lipids and as class averages. These changes encompassed ceramides, PS, PE and PC phospholipids and SM lipids. As these changes could arise from a variety of altered enzymatic pathways, we sought to use isotope tracing flux analysis to quantify phospholipid production fluxes and highlight altered pathways. We expanded deuterium tracer methodology previously used in cell lines, healthy animals, and human volunteers (Bleijerveld et al., 2007; DeLong et al., 1999; Peng et al., 2021; Pynn et al., 2011) to quantify the production fluxes of PC and PE lipids in zebrafish tumors.

Bavituximab, a monoclonal antibody against PS lipids, showed promising results in combination with sorafenib in mouse xenografts and is now in early clinical trials (Cheng et al., 2016). Both xenograft models presented in this study had activating mutations in Wnt/β-catenin signaling pathway components. Our finding that activated β-catenin enhanced synthesis fluxes of PS lipids in human liver cells provides further rationale for targeting phospholipid metabolism in HCC, especially those with mutations in *CTNNB1* or other Wnt/β-catenin signaling pathway genes.

While we didn’t observe a significant decrease in total PC lipids in our experiments, other groups have recently reported a significant reduction in total PC lipid abundances in HCC from human patients (Ismail et al., 2020; Krautbauer et al., 2016). Our observation of decreased rates of total PC lipid synthesis in β-catenin-driven tumors should lead to a reduction in total liver PC lipids and is a possible mechanism that supports these observations. Our inability to observe changes in total PC lipid abundances in zebrafish could be due to reduced PC lipid export in zebrafish relative to humans. A long-standing practice to accelerate HCC growth in mouse models involves limiting choline and methionine in the diet (Haberl et al., 2020; Ikawa-Yoshida et al., 2017; Yoshida et al., 1993), which is thought to suppress *de novo* PC lipid synthesis and increase diet-induced lipotoxicity. Conversely, choline supplementation blunts the development of carcinogen-induced HCC(Brown et al., 2020). Our data suggest that oncogenic mutations in *CTNNB1* might drive hepatocarcinogenesis through a similar mechanism via decreased PC synthesis.

In zebrafish HCC, we identified large changes in the lipidome that clearly distinguished male from female livers. These sex-specific differences in metabolism were further highlighted by phospholipid flux analysis. We observed large difference in PC synthesis fluxes between male and female fish when we analyzed 9-^2^H choline and 4-^2^H ethanolamine tracing data. Male and female Tg-ABC zebrafish had sex-dependent mechanisms to achieve similar ends: 9-^2^H choline highlighted decreased PC lipid synthesis through the CDP-choline pathway in female, but not in male, Tg-ABC zebrafish; conversely, 4-^2^H ethanolamine identified decreased PC synthesis from PE methylation in male, but not in female, Tg-ABC zebrafish (Fig 6E,F). Similar to male zebrafish, *4x-Ala-CTNNB1* THLE-2 cells, which were derived from a male patient (Pfeifer et al., 1993), did not show conversion of PE lipids into PC lipids.

The finding of sex-based differences in PC lipid metabolism in HCC underscores the importance of sex as a biological variable in hepatocarcinogenesis and lipid metabolism. In humans, it has been reported that PEMT activity is increased by estrogen, and therefore females may be resistant to choline deprivation (Leermakers et al., 2015). Sex-specific differences have been well characterized in both HCC (Natri et al., 2019; Takemoto et al., 2005) and MASLD, MASH, and HCC risk (Lonardo et al., 2019), as well as in mouse primary hepatocytes (Hochmuth et al., 2021), obese mice (Savva et al., 2022), rats (Lomas-Soria et al., 2018), and the zebrafish liver transcriptome (Zheng et al., 2013). At least some of the increased susceptibility to hepatocarcinogenesis in males is due to the effects of estrogen, which inhibits pro-tumorigenic IL-6 production (Naugler et al., 2007). Our findings motivate further research into sex-specific differences in HCC lipid metabolism, which may influence response to systemic therapies.

## Materials and Methods

### Animal Husbandry

Zebrafish (*Danio rerio*) lines were maintained at the University of Utah core zebrafish facilities under standard conditions in compliance with the University of Utah Institutional Animal Care and Use Committee guidelines (Kimmel et al., 1995). Transgenic zebrafish expressing hepatocyte-specific activated β-catenin, *Tg(fabp10a:pt-*β*-cat)* (Evason et al., 2015), and their non-transgenic siblings were used for this study. Embryos and larvae were cultured in egg water (2.33 g Instant Ocean in 1 liter of Milli-Q water with 0.5 mL Methylene Blue), incubated at 28.5°C, sorted by eye color at 3-5 dpf (with green eyes indicating ABC transgenic fish due to *cryaa:venus*), and raised to at least 4 months of age. Adult zebrafish were fed brine shrimp, flakes, and powdered food and housed on a recirculating system.

### Cell lines

The immortalized human liver epithelial cell line THLE-2 was purchased from American Type Culture Collection (ATCC, CRL-2706). THLE-2 cells were maintained on plates coated with 10% filter-sterilized collagen from calf skin (Sigma Aldrich, C8919) at 37°C and 5% CO_2_. Cells were cultured using BEBM Basal Media (Lonza Bioscience, CC-3171) supplemented with the BEGM Bronchial Epithelial SingleQuots™ Kit (Lonza, CC-4175), excluding epinephrine and GA-1000. The media was supplemented with 10% fetal bovine serum (FBS) (Thermo, 10437028) and 100 U/mL penicillin/100 µg/mL streptomycin (Thermo, 15140122).

### Lentivirus Production

We used the Lenti-X™ Tet-On® Advanced Inducible Expression System (Clontech, 632162) to produce the lentivirus vectors and transfect cells. We first prepared three lentivirus vectors that express our target genes, namely the pLVX-Tet-On Advanced Vector and the pLVX-Tight-Puro Vector containing either a WT *CTNNB1* construct (pLVX-Tight-Puro-WT-CTNNB1) or mutant *CTNNB1* (pLVX-Tight-Puro-4x-Ala-CTNNB1), an allele containing mutations (S33A, S37A, T41A, S45A). Following vector preparation, 293T/17 cells (ATCC, CRL-11268) were seeded on 10 cm tissue culture plates in DMEM (Thermo, 11965092) at 3.0×10^6^ cells per plate. The plates were then incubated at 37 °C and 5% CO_2_ for 24 hrs. Subsequently, 293T cells were transfected with each of the above-mentioned vectors and Lenti-X Packaging Single Shots (VSV-G) (Clontech, 631275) in the presence of reduced-serum Opti-MEM medium (Gibco, 31985062) and 1 mg/mL polyethylenimine (PEI) (Polysciences, 23966-1) in a 4:1 ratio of DNA (μg):PEI (μg), and medium in a 1:9 ratio of DNA/VSV-G/PEI mixed solution (mL):medium (mL). The transfected cells were then incubated at 37 °C and 5% CO_2_ for 24 hrs. The medium was removed and replaced with 4 mL of growth medium. The cells were then incubated at 37°C and 5% CO_2_ for 6 hrs, followed by the collection and filtration of the medium through a 0.45 μm PVDF filter. 5 mL of the medium was subsequently replaced, and cells were incubated at 37 °C and 5% CO_2_ for 12 hrs, followed by the collection and filtration of the medium as described above. The collected medium was stored at 4 °C until further use. The stored medium was ultracentrifuged at 4 °C and 112,000 × g for 105 min to produce high-titer lentivirus vectors. The solution was stored at −80 °C in cryovials containing 2 mL each for subsequent lentivirus infection.

### Lentivirus Transfection and Doxycycline Induction

THLE-2 cells were seeded onto T-75 flasks containing growth medium at 2.0×10^6^ cells per plate. The growth medium was removed after two days and cells were infected with 2 mL of either the pLVX-Tight-Puro-WT-CTNNB1 lentivirus or the pLVX-Tight-Puro-4xAla-CTNNB1 lentivirus as well as 2 mL of medium containing the pLVX-Tet-On lentivirus. This was followed by treatment with polybrene (Santa Cruz Biotechnology, sc-134220) to a final concentration of 3 μg/mL. Flasks were gently rocked every 30 min for 6 hrs, followed by the addition of 2 mL of fresh medium and incubation at 37 °C and 5% CO_2_ for 24 hrs. Subsequently, another 10 mL of fresh medium was added, followed by polybrene treatment to a final concentration of 3 μg/mL. The cells were incubated at 37 °C and 5% CO_2_ for 24 hrs and selected with growth medium containing 6 μg/mL puromycin (Sigma-Aldrich, P8833) and 750 μg/mL neomycin (Thermo, 10131035) for 48 hrs. The clones were initially treated with 0.5 μg/mL doxycycline (Sigma-Aldrich, D5207) for 48 hrs to induce the expression of target genes. 0.25 µg/mL of puromycin and 100 µg/mL of neomycin were used to maintain the doubly transfected cell lines. To maintain doxycycline-mediated induction of β-catenin, cells were kept in 1 µg/mL of filter-sterilized dox (D5207).

### Cell Lysate and Western Blotting

THLE-2 cells were plated at a density of 5.0×10^5^ cells/mL on collagenated 6 cm plates and allowed to grow for 24 hrs at 37°C and 5% CO_2_. The cells were washed with cold PBS and lysed in RIPA buffer (50 mM Tris, 150 mM NaCl, 0.1% SDS, 0.5% sodium deoxycholate, 1% NP-40) supplemented with 1 mM phenylmethylsulfonyl fluoride (PMSF), 1 mM sodium orthovanadate, and 1% protease inhibitor cocktail (Sigma, P2714). Protein concentration was quantified using the Pierce BCA Protein Assay Kit (Thermo, 23225). Samples were mixed with 4X Laemmli Sample Buffer (BioRad, 1610747) and 2-mercaptoethanol to a final concentration of 2.5% and incubated at 90°C for 5 min. 10 μg of total protein was resolved on a 4-20% polyacrylamide gel (BioRad, 4561096) at 80 V for 30 min then 100 V for 45 min. Gels were blotted on 0.2 µm nitrocellulose membranes (BioRad, 1704270) via the Trans-Blot Turbo Transfer System (BioRad, 1704150) according to standard procedure at 25 V for 30 min. After blocking with 5% non-fat milk (Sigma, M7409)/Tris-buffered saline with 0.05% Tween 20 (TBS-T) for 1 hr, the membrane was washed with TBS-T and incubated overnight in 5% bovine serum albumin (Sigma, A6003)/TBS-T with 1:1000 anti-β-Catenin primary antibody (Cell Signaling, 8480). The membrane was then washed with TBS-T and incubated with 1:10000 goat anti-rabbit poly-HRP secondary antibody (Thermo, 32260) and 1:10,000 anti-β-actin-peroxidase antibody (Sigma, A3854) in 5% bovine serum album/TBS-T for 1 hr. The membrane was then washed with TBS-T and chemiluminescence was assessed with the BioRad ChemiDoc MP Imaging System (BioRad, 12003154).

### RT-qPCR

THLE-2 cells were plated at a density of 5.0×10^5^ cells/mL on collagenated 6-well plates and allowed to grow for two days at 37°C and 5% CO_2_. Cells were washed with PBS and RNA was isolated using the Qiagen RNeasy Mini Kit (Qiagen, 74104) according to the manufacturer’s specifications. cDNA was then generated using QuantaBio qScript cDNA SuperMix (QuantaBio 101414-102). qPCR master mixes were prepared consisting of 2.5% 100 µM forward primer, 2.5% 100 µM reverse primer, and 62.5% PowerTrack SYBR Green Master Mix (Thermo, A46109) in RNase-free water. Master mixes were combined 4:1 with the cDNA reactions and plated in duplicate. qPCR was performed using the LC480 PCR Lightcycler (Roche, 05015278001) using the “Mono Color Hydrolysis Probe/UPL probe” detection format. The temperature cycle consisted of an initial 2 min period at 95°C and 40 cycles of 95°C for 15 sec and 60°C for 50 sec set to single acquisition mode. The housekeeping gene *RPS2* was used as an internal control for cDNA quantification and normalization of the amplified products. All data are reported as mean ± stdev.

### Anchorage-Independent Growth Assay

Matrigel (Corning 354234) was used to coat the bottom of 48-well plates and allowed to solidify at 37°C for 30 min. All plates, pipette tips, and Matrigel were kept on ice to prevent spontaneous solidification of the Matrigel. Pelleted THLE-2 cells were diluted to 2.0 × 10^5^ cells/mL in media and combined 1:1 with media containing 4% Matrigel followed by incubation at 37°C for 7 min. 1 µg/mL dox and 10 µM myriocin were added to the relevant samples at this time. The cell suspension was plated above the solidified Matrigel and allowed to grow for four days at 37°C and 5% CO_2_. 0.3 mg/mL resazurin sodium salt (Sigma, R7017) in media was added to the cell suspensions to a final volume of 10% and allowed to incubate at 37°C and 5% CO_2_ for 2 hrs. Fluorescence was measured on the Biotek Neo2 Plate Reader (Agilent) with excitation/emission of 560 nm/590 nm. Readings were normalized by subtracting the fluorescence value from a no cell control well. All data are reported as mean ± stdev.

### Zebrafish Metabolic Profiling Preparation

Male and female *Tg(fabp10a:pt-*β*-cat)* zebrafish were raised to 5 months post-fertilization (mpf) under standard conditions. Animals were fasted for 12 hrs before euthanasia. Livers were isolated using a dissecting microscope and weighed. For non-transgenic samples, liver tissues from up to four animals of the same sex were pooled to achieve a minimum of 20 mg. Livers were snap frozen in liquid nitrogen and placed on dry ice and submitted for LC-MS and GC-MS analysis (see below).

### Zebrafish Isotope Tracing

Zebrafish IP injection was performed using standard methods (Kinkel et al., 2010). In brief, zebrafish were anesthetized with tricaine and weighed. Zebrafish IP injection was performed with 1 μL of solution per 100 mg of body weight using a 10 μL NanoFil syringe with a 35G beveled steel needle (World Precision Instruments). Injection solutions consisted of 2 mg/mL 9-^2^H choline, 2 mg/mL 4-^2^H ethanolamine, 30 mg/ml ^13^C glucose or 5 mg/mL ^13^C glutamine dissolved in 0.9% sterile saline (SteriCare Solutions, 6281) prepared immediately prior to injection. Under a dissection scope, anesthetized zebrafish were placed on a sponge wetted with tricaine system water and gently held abdomen up. The needle was inserted into the midline between the pelvic fins and the total volume was injected by fully depressing the plunger in a swift and controlled movement. After injection, zebrafish were placed into a static tank with fresh system water and checked for injury and bleeding. Resuscitation was immediately performed by using a transfer pipette to move water across the gills until there was spontaneous muscle movement and the zebrafish was able to remain upright. After intraperitoneal (IP) injection, zebrafish were stored in static housing for 24 hrs, without scheduled feedings. Wellness checks were performed at 3- and 18-hours post exposure, and any injured zebrafish were immediately euthanized. At the desired endpoint, injected zebrafish were euthanized and, using a dissection microscope, livers were dissected, weighed, and immediately placed on dry ice. The genotype for each euthanized zebrafish was confirmed using eye color under a fluorescent microscope.

### TCGA and Zebrafish Transcriptomics Analysis

Human patient paired tumor and non-tumor HTSeq transcriptomic data was gathered from The Cancer Genome Atlas Liver Hepatocellular Carcinoma (TCGA-LIHC) GDC 23.0 Data Release using TCGAbiolinks package (Colaprico et al., 2016). Patient samples were excluded from analysis if history indicated they had received treatment prior to biopsy or had a final diagnosis of a malignancy other than, or in addition to, HCC. In total, 45 patients were eligible for analysis. The raw counts were normalized and differentially expressed genes were identified using a 5% false discovery rate with DESeq2 version 1.34.0 (Love et al., 2014). To perform a nested analysis comparing patients with and without mutations in CTNNB1, hereto referred as LIHC-Nested, mutation status was determined by MuSE v2.0 (Fan et al., 2016). Overall, 12 out of 45 patients had mutations in *CTNNB1*, with eleven men and one woman. To identify specific individual events, 11 male patients with non-mutated *CTNNB1* were randomly selected as controls for the 11 male patients with mutated *CTNNB1* and the lone woman patient was excluded. The tumor and control samples were first compared in *CTNNB1*-mutated samples while controlling for individual effects and then in the CTNNB1-non-mutated samples. *CTNNB1*-mutated and non-mutated samples were then combined into a single model to test interactions and main effects by following the group-specific condition effects, individuals nested within groups section of the DESeq2 vignette.

Transcriptomics data from male (Kalasekar et al., 2019) and female (Evason et al., 2015) zebrafish were re-analyzed from previously published datasets. Raw counts were normalized and differentially-expressed genes were identified as stated above with DESeq2 version 1.34.0, using GRCz11 genome build. Each dataset contained RNASeq data for 5 transgenic ABC zebrafish and 5 non-transgenic, sex-matched, control siblings.

Given the small sample size of each zebrafish dataset and heterogenous nature of the TCGA dataset, genes were considered significantly dysregulated if L2FC was greater than +/− 0.5 with a padj<0.05. GeneOntology was used for GO enrichment analysis, by comparing biological process for genes with a padj<0.05 and L2FC greater than +/− 1.0 for each male zebrafish, female zebrafish, TCGA-LIHC, and LIHC-Nested. GO enrichment analyses were then compared between sets for identical and related GO terms.

### *In vitro* sample preparation for LC-MS

THLE-2 cells were plated on collagenated 10 cm plates at a density of 7.0×10^5^ cells per plate using media supplemented with 10% dialyzed FBS and no penicillin/streptomycin. Cells were allowed to grow at 37°C and 5% CO_2_. For untargeted quantitative lipidomics, the cells were cultured for 48 uninterrupted hours. For stable isotope tracing, labeled and unlabeled metabolites were added to the respective experimental and control plates after 24 hrs at a final concentration of 250 µM for ethanolamine (4-^2^H ethanolamine), 400 µM for serine (3-^2^H serine), and 100 µM for choline (9-^2^H choline), followed by an additional 24 hrs of growth. A designated plate was used to determine the total cell volume of each plate. Experimental plates were washed once with PBS and then pelleted in cold PBS at 200 x g for 5 min. A lipid extraction solution consisting of 75% methyl *tert*-butyl ether, 23% methanol, 1% Splash Lipidomix Mass Spec Standard (Avanti Polar Lipids, 330707), and 1% Deuterated Ceramide Lipidomix Mass Spec Standard (Avanti Polar Lipids, 330713) was created and 1 mL was used to resuspend each cell pellet. The extraction mixture was allowed to rest on ice for 15 min with occasional vortexing. 20% water was added to induce phase separation of the mixture followed by centrifugation at 4°C for 10 min. The organic layer was transferred to a secondary glass container while the aqueous layer was recentrifuged at max speed and 4°C for 10 min followed by transfer of the supernatant to a separate glass container. The organic and aqueous fractions were dried gaseous nitrogen. The organic samples were resuspended in 35x the packed cell volume using 2:1:1 isopropyl alcohol:acetonitrile:water and transferred to amber glass mass spectrometry vials (Agilent, 5182-0716) containing a glass insert. The aqueous samples were resuspended in 35x the packed cell volume using 80:20 methanol:water and transferred to polypropylene mass spectrometry vials (Agilent, 5188-2788).

### *In vivo* sample preparation for LC-MS

20-30 mg sections of snap frozen zebrafish liver tissue were transferred to pre-chilled Safe-Lock tubes (Eppendorf, 022363352) containing a cold 5/16 in. diameter stainless steel ball (Grainger, 4RJL8). The tissue was disrupted by shaking at 25 Hz for 30 sec under liquid nitrogen using the Retsch CryoMill (Retsch, 20.749.0001). A lipid extraction solution consisting of 76% methyl *tert*-butyl ether, 23% methanol, and 1% Splash Lipidomix Mass Spec Standard (Avanti Polar Lipids 330707) was added to the homogenized tissue.

Metabolite extraction was performed identically to the *in vitro* sample preparation method described above. The organic samples were resuspended in 200 µL of 2:1:1 isopropyl alcohol:acetonitrile:water and transferred to amber glass mass spectrometry vials (Agilent 5182-0716) containing a glass insert. The aqueous samples were resuspended in 35X the tissue mass (mg) of 80:20 methanol:water and transferred to polypropylene mass spectrometry vials (Agilent 5188-2788).

### LC-MS Methodology

Extracted aqueous and polar metabolites were analyzed by LC-MS using a Vanquish HPLC system (Thermo Fisher Scientific) and a QExactive HF Orbitrap mass spectrometer (Thermo Fisher Scientific). For hydrophobic metabolites (lipids), separation was achieved by C18 chromatography performed on an Acquity UPLC CSH C18 column (2.1 mm X 100 mm, 1.7 µm particular size, 130 Å pore size, Waters Co., 186005297). The chromatography gradient was formed by solvent A (10 mM ammonium formate and 0.1% formic acid in 60:40 acetonitrile:water) and solvent B (10 mM ammonium formate and 0.1% formic acid in 90:9:1 isopropyl alcohol:acetonitrile:water) at a constant flow rate of 350 µL/min. The gradient function was: 0 min, 30% B; 5 min, 43% B; 5.1 min, 50% B; 14 min, 70% B; 21 min, 99% B; 24 min, 99% B; 24.1 min, 30% B; 31 min, 30% B. For aqueous phase polar metabolites, separation was achieved by hydrophilic interaction liquid chromatography (HILIC) performed on an Atlantis Premier BEH Z-HILIC column (2.1 mm X 50 mm, 1.7 µM particular size, 95 Å pore size, Waters Co., 186009978) run with a gradient of solvent A (10 mM ammonium acetate in 100% water, pH 9.2) and solvent B (100% acetonitrile) at a constant flow rate of 350 µL/min. The gradient function was: 0 min, 95% B; 10.4 min, 45% B; 11.5 min, 45% B; 11.6 min, 95% B; 15 min, 95% B. Autosampler temperature was 4°C, column temperature was 30°C, and injection volume was 2 µL. Samples were injected into the mass spectrometer by electrospray ionization operating in positive ion mode. Lipid samples were analyzed using a ddMS2 method with a resolving power of 35,000 at m/z of 200 and loop count 15 / isolation window 1.0 m/z / (N)CE/stepped (N)CE 25,30 as well as a full scan method with a resolving power of 70,000 at m/z of 200 and range of 150 – 1500 m/z. Aqueous samples were analyzed using a full scan method with a resolving power of 70,000 at m/z of 200 and range of 74 – 1110 m/z. Full scan data were analyzed using the Maven software package with specific peaks assigned based on exact mass and comparison with known standards (Clasquin et al., 2012). Extracted peak intensities were corrected for natural isotopic abundance using the R package AccuCor (Su et al., 2017) in R version 4.2.2.

### Quantitative lipidomics analyses

ddMS2 data were analyzed using the LipidSearch 5 software package (Thermo) with LC-MS search performed using standard parameters and LC-MS alignment performed using a median calculation method and otherwise standard settings. Mean peak areas for all identified lipid species were taken from the LipidSearch 5 program. This dataset was supplemented by a manual peak assignment using Maven for standards and key lipid species not identified by LipidSearch. Lipid species with at least 1 missing value were removed from all *in vitro* datasets while, due to the larger sample size, lipid species with at least 3 missing values were removed from all *in vivo* datasets. All lipids possessing a median peak area < 1000 were removed. All PC and PE lipids with median peak area < 100,000 were dropped, resulting in the exclusion of the approximately 20% least abundant lipids in these classes. The peak areas were normalized by lipid class to the relevant standard molecule in the applied standard mixes. For lipid classes without a directly comparable standard, the average area across all standards was used for normalization. Volcano plots were generated using the R package EnhancedVolcano (Blighe et al., 2023). Principal component analysis was performed using MetaboAnalyst 5.0 (Pang et al., 2021) with no data filtering, log base 10 transformation, and pareto scaling.

### Quantitative isotope enrichment analyses

For lipid isotope enrichment analysis, lipids mass spectra were collected in full scan mode at 70,000 resolving power on an Orbitrap QE plus mass spectrometer run over a C18 column using the gradient function described above. Lipid peak ID’s were based on exact mass and retention times determined by ddMS2 analysis for a subset of high confidence phospholipid species with raw ion intensities of greater than 1e7 that were well separated from isomeric peaks. Isotopic peaks were picked using MAVEN. The labeling fraction (*frac_lab_*) of a lipid is determined from ^2^H natural isotope abundance by the following equation:

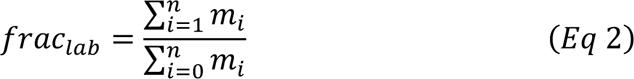

Where *n* is the number of heavy labeled atoms in the molecule, *i* is each mass isotopologue (m+0, m+1 etc.) and *m* is the corrected abundance of mass isotopologue *i*. For cell lines, tracer enrichment was determined by isotopic fractional enrichment of the tracer at the end time point. For zebrafish, a separate cohort of fish were injected with tracer and euthanized at 2, 8 and 24 hours post injection to determine tracer exposure kinetics. Enrichment fractions were fit to a decay function and total exposure determined by integration of that function over the total exposure time (24 hrs). From fractional labeling data, FSR (hr-1) was calculated for each lipid species (equation 1). Absolute synthesis rate (*R_syn_*) or flux was solved for by multiplying FSR by the measured abundance of the lipid:

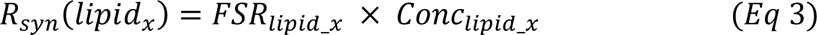

Where *Conc* is the concentration of lipid_x (pmoles/mg tissue).

### Zebrafish GC-MS

Metabolites were extracted from frozen tissue in 225 µL ice-cold MeOH containing internal standards (d4-succinate, 1 µg/sample; d9-carnitine, 1 µg/sample), 750 µL of ice-cold MTBE (methyl tert-butyl ether) and 400 µL of water in bead mill tubes (0.4 mm ceramic). The sample was homogenized in one 30 s cycle followed by centrifugation at 20,000 g for 5 min at 4 °C. The lower phase containing the metabolite fraction was collected and evaporated to dryness under vacuum. A pooled QC sample was prepared by taking 10 µL aliquots from each sample prior to dry down.

All GC-MS analysis was performed with a Waters GCT Premier mass spectrometer fitted with an Agilent 6890 gas chromatograph and a Gerstel MPS2 autosampler. Dried samples were suspended in 40 µL of a 40 mg/mL O-methoxylamine hydrochloride (MOX) in pyridine and incubated for one hour at 30°C. To autosampler vials was added 25 uL of this solution. 40 µL of N-methyl-N-trimethylsilyltrifluoracetamide (MSTFA) was added automatically via the autosampler and incubated for 60 minutes at 37°C with shaking. After incubation 3 µL of a fatty acid methyl ester standard (FAMES) solution was added via the autosampler then 1 uL of the prepared sample was injected to the gas chromatograph inlet in the split mode with the inlet temperature held at 250°C. A 10:1 split ratio was used for analysis. The gas chromatograph had an initial temperature of 95°C for one minute followed by a 40°C/min ramp to 110°C and a hold time of 2 minutes. This was followed by a second 5°C/min ramp to 250°C, a third ramp to 350°C, then a final hold time of 3 minutes. A 30 m Phenomenex ZB5-5 MSi column with a 5 m long guard column was employed for chromatographic separation. Helium was used as the carrier gas at 1 mL/min. Due to the high amounts of several metabolites the samples were analyzed once more at a 10-fold dilution.

## Data Transparency

All data and analysis will be made freely available to the research community.

## Acknowledgements

We thank all members of the Ducker and Evason labs for their contributions to this research. We acknowledge Dr. Mei Yee Koh for technical assistance. We thank Alan Maschek for technical assistance in lipid mass spectrometry methods. This research was supported in part by funding to G.S.D. and K.J.E. from the Damon Runyon Cancer Research Foundation (Damon Runyon-Rachleff Innovation Award DR 61-20), funding to G.S.D from the National Cancer Institute (R00CA215307) and the American Cancer Society (DBG-23-1037804-01-TBE) and the University of Utah School of Medicine, and funding to K.J.E. from the National Cancer Institute (R01CA222570) and the American Cancer Society (RSG-22-014-01-CCB). GCMS Metabolomics analysis was performed by the Metabolomics Core at the University of Utah Health Sciences Centers using equipment provided for by University of Utah RIF funds.

## Author Contributions

G.S.D. and K.J.E. designed the study. C.V.W., H.L., J.K., C.S., T.Q., A.S., A.R., C.C. performed experiments. C.V.W., H.L., T.Q. and G.S.D. performed data and statistical analysis. G.S.D, K.J.E., S.A.S, C.V.W. and J.E.C. interpreted the data. C.V.W., H.L., K.J.E. and G.S.D. wrote the manuscript. All authors had access to the study data and had reviewed and approved the final manuscript.

## Supplemental Files

Supplemental Data File 1: Quantified lipid abundances and turnover rates from 4x-Ala-CTNNB1 and WT-CTNNB1 THLE-2 cells

Supplemental Data File 2: Quantified lipid abundances and turnover rates from non-Tg and Tg-ABC zebrafish livers

Supplemental Data File 3: Individual GeneOntology Analyses

Supplemental Data File 4: Collated Transcriptomic Analysis of Metabolic Pathways in Human and Zebrafish ABC-driven HCC.

Supplemental Data File 5: Primers used for qPCR quantification

---

**Supplemental Figure 1:**
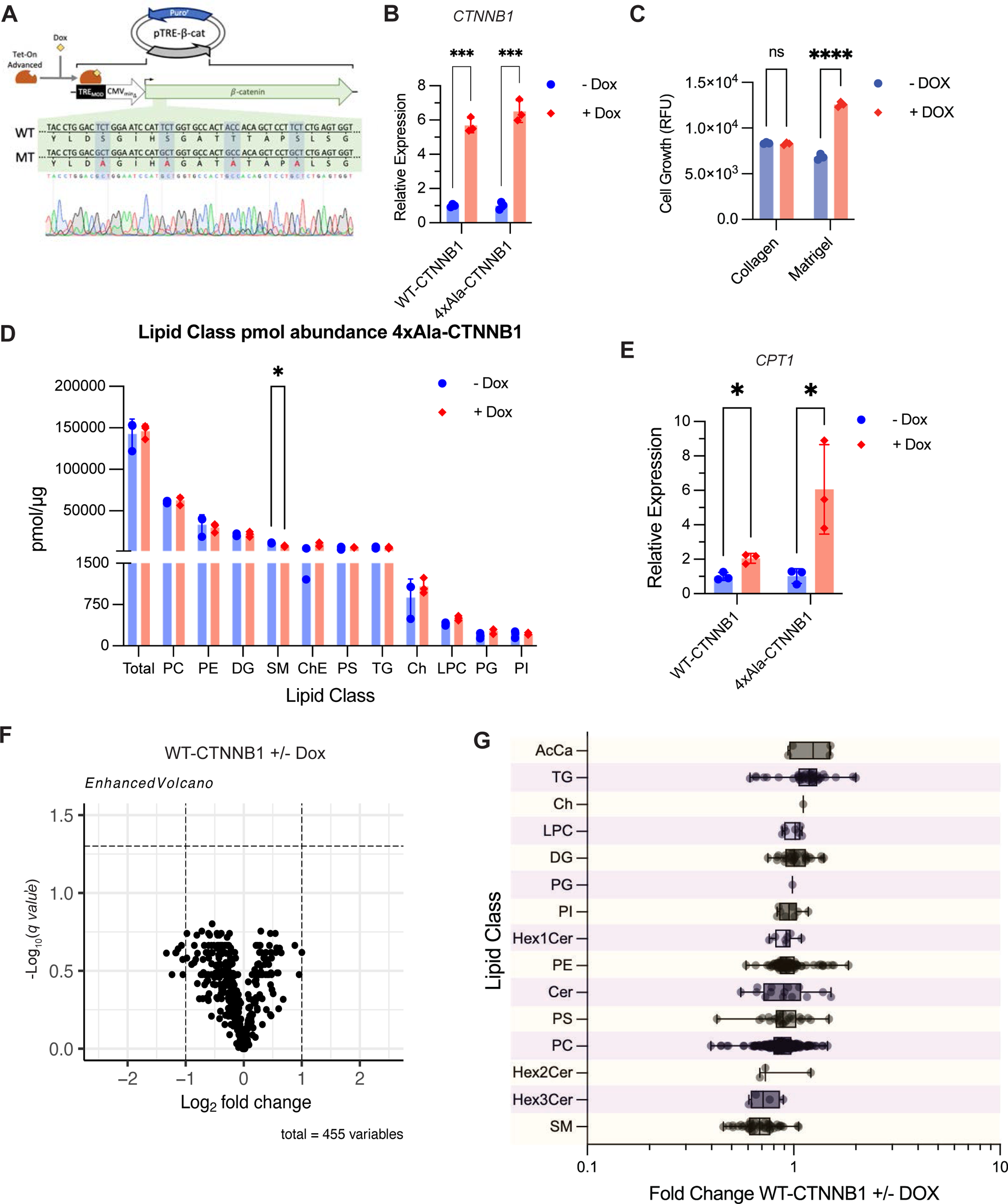
Verification of *CTNNB1* overexpression constructs and activated β-catenin-dependent effects on lipid metabolism. A. Overview of Tet-On Dox-inducible β-catenin system showing WT and quadruple alanine mutant (MT) *CTNNB1* constructs. Sequencing verification for the *4x-Ala-CTNNB1* mutant. B. qRT-PCR of the β-catenin gene *CTNNB1* from THLE-2 cells +/− Dox treatment in *CTNNB1* overexpression constructs (n=3, mean±stdev). C. Growth assay of THLE-2 cells +/− Dox treatment in *4x-Ala-CTNNB1* overexpression construct grown under adherent (collagen) or anchorage independent (Matrigel) conditions (n=3, mean±stdev). D. Total quantified abundance of different lipid classes from THLE-2 cells +/− Dox treatment in *4x-Ala-CTNNB1* overexpression construct (n=3, mean±stdev). E. qRT-PCR of the rate-limiting enzyme in fatty acid oxidation *CPT1* from THLE-2 cells +/− Dox treatment in *CTNNB1* overexpression constructs (n=3, mean±stdev). F. Volcano plot showing change in abundance for individual lipid species from THLE-2 cells +/− Dox treatment in *WT-CTNNB1* overexpression construct (n=3). Y-axis is the −Log_10_q using a multiple-comparisons corrected false-discovery rate of 5%. G. Fold changes in lipid species by class. Lipid classes are defined in Fig. 1. Fold change is average change between *WT-CTNNB1* expressing THLE-2 cells +/− Dox (n=3). RFU = relative fluorescence units. For B, C, D, and G, * p<.05, ** p<.01, *** p<.001, **** p<.0001 by multiple unpaired t-tests, corrected for multiple comparisons using the Holm-Šídák method.

**Supplemental Figure 2:**
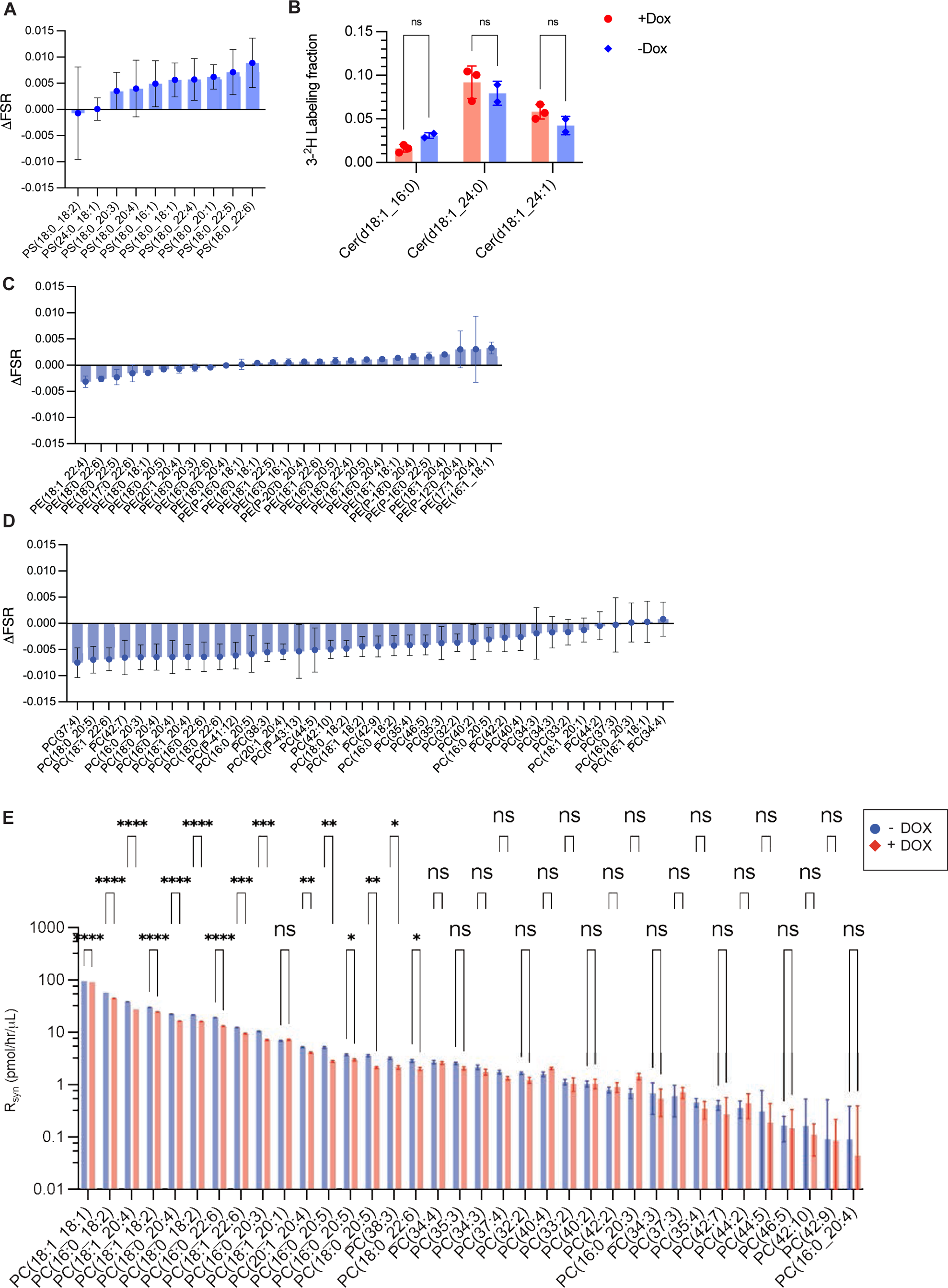
Activated β-catenin drives directional alterations in phospholipid synthesis rates. A. Change in FSR of PS lipids for THLE-2 cells +/− Dox in *4x-Ala-CTNNB1* overexpression construct. Difference reflects FSR with Dox induction subtracted by FSR without Dox induction (n=3, error bars reflect propagation of error from FSR calculations). B. 3-2H serine incorporation into ceramides in THLE cells (n=3, mean±stdev, n.s. by unpaired t-test). C. Change in FSR of PE lipids for THLE-2 cells +/− Dox in *4x-Ala-CTNNB1* overexpression construct (n=3, error bars reflect propagation of error from FSR calculations). D. Change in FSR of PC lipids for THLE-2 cells +/− Dox in *4x-Ala-CTNNB1* overexpression construct (n=3, error bars reflect propagation of error from FSR calculations). E. R_syn_ of PC lipids from THLE-2 cells +/− Dox in *4x-Ala-CTNNB1* overexpression construct (mean±stdev). For E, * p<.05, ** p<.01, *** p<.001, **** p<.0001 by multiple unpaired t-tests, corrected for multiple comparisons using the Holm-Šídák method.

**Supplemental Figure 3:**
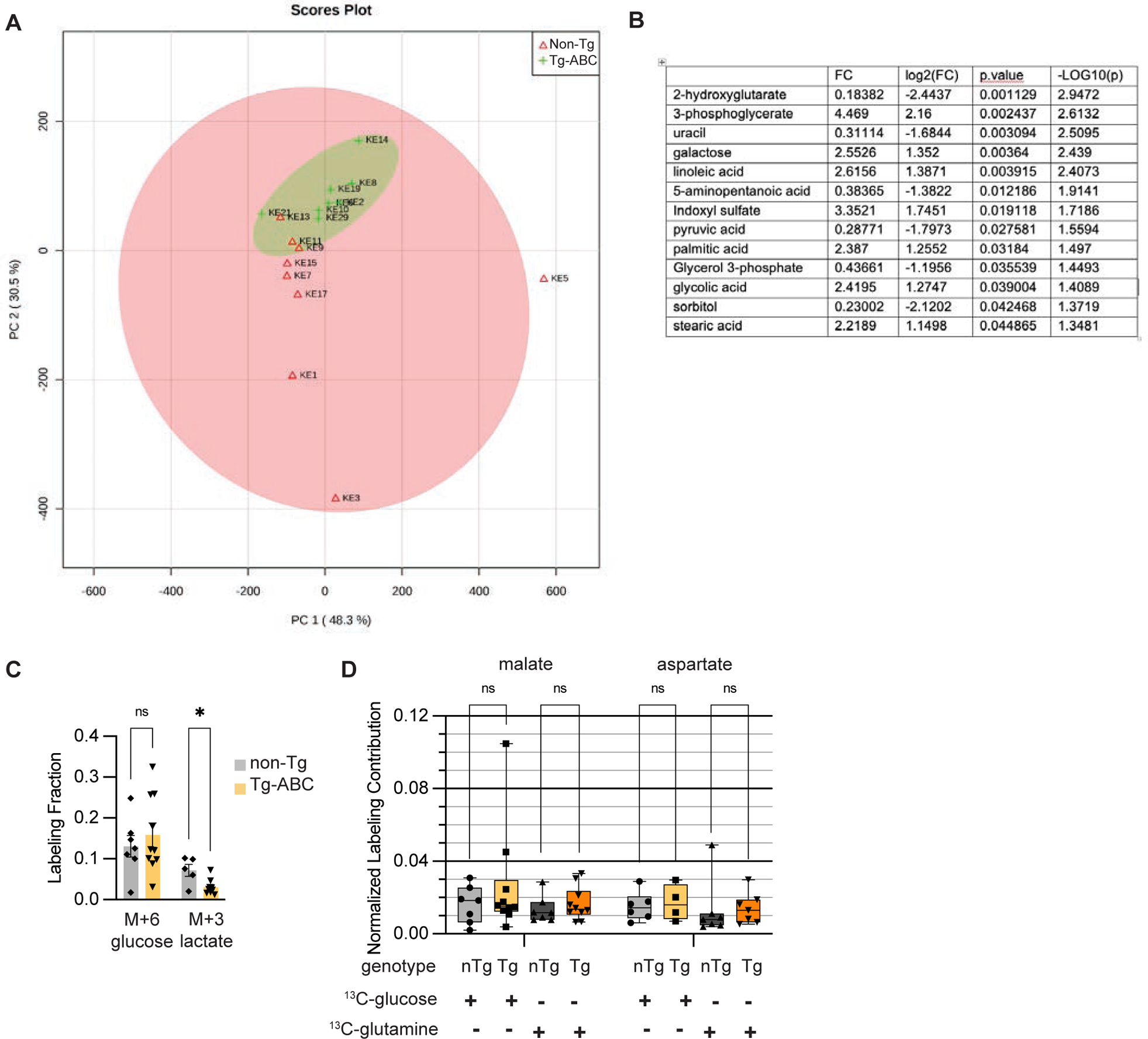
Tg-ABC zebrafish show a notable decrease in glycolytic metabolism. A. Principal component analysis of all quantified polar metabolites extracted from livers of non-Tg and Tg-ABC zebrafish (n=8-9). B. Table showing average fold change (FC) in abundance for significant polar metabolites from non-Tg and Tg zebrafish livers (n=8-9). A significance cut-off of at least magnitude 1 log base 2 fold change (log2(FC)) and raw p-value of at least 0.05 was used. C. 6-^13^C labeling fraction of glucose and 3-^13^C labeling fraction of lactate in non-Tg and Tg-ABC zebrafish livers 2 hrs after injection with 6-^13^C glucose (n=5-10, mean±SEM). D. Normalized isotopic labeling contributions to key TCA intermediates 2 hrs after injection with either 6-^13^C glucose or 5-^13^C glutamine in non-Tg (nTg) and Tg-ABC (Tg) zebrafish livers (n=4-10, mean±SEM). For C-D, * p<.05 by multiple unpaired t-tests, corrected for multiple comparisons using the Holm-Šídák method.

**Supplemental Figure 5:**
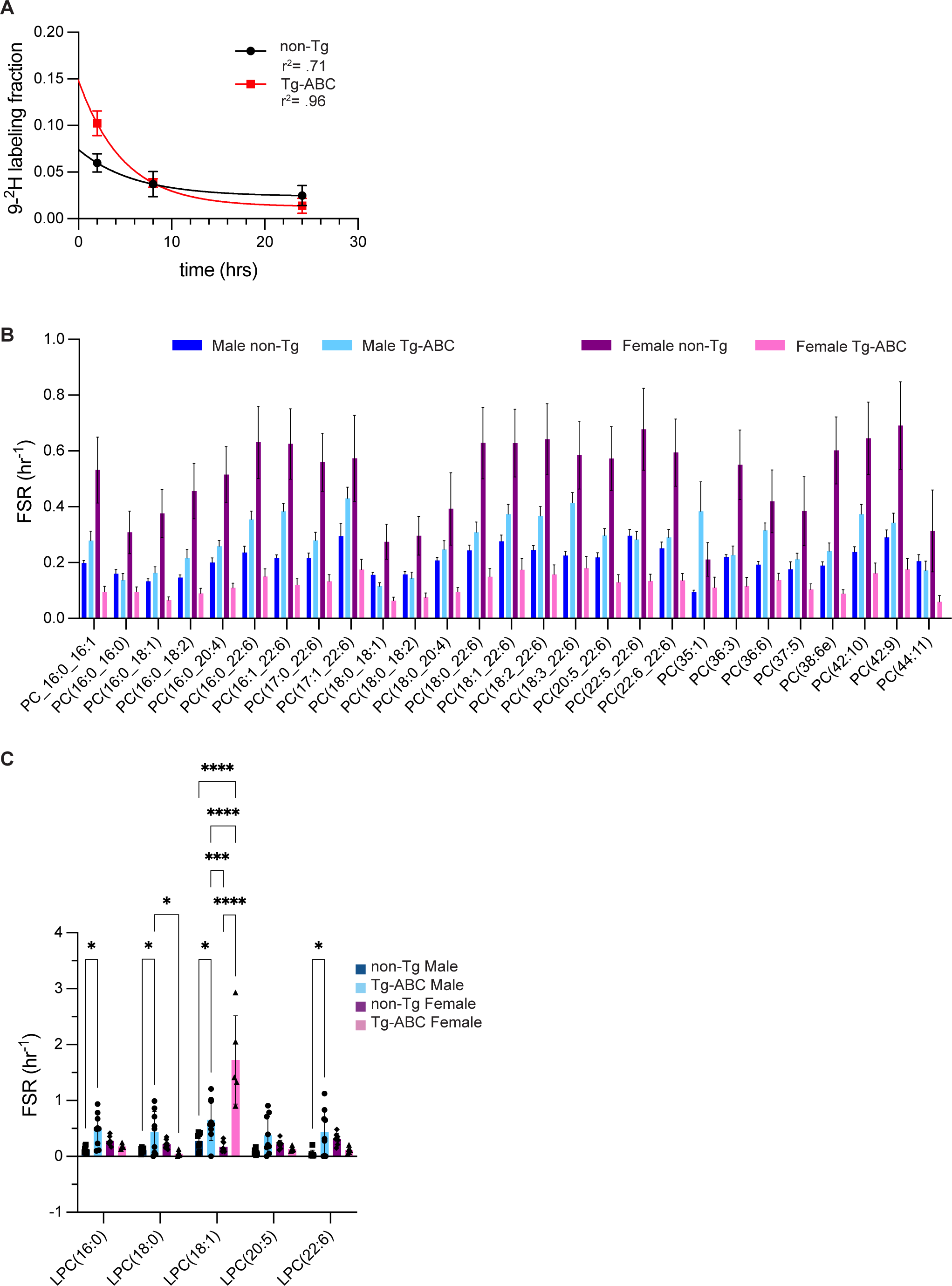
Sex and genotype-specific differences in PC and LPC synthesis rates using a labeled choline tracer. A. Recovered labeling fraction of phosphocholine in non-Tg and Tg-ABC zebrafish livers 2 hrs, 8hrs, and 24 hrs after 9-^2^H choline injection, fit to first-order decay kinetics (n=2-4, mean±SEM). B. FSR of PC lipids as calculated from 9-^2^H choline incorporation in non-Tg and Tg-ABC, male and female zebrafish livers 24 hrs after injection (n=5-9, mean±SEM). C. FSR of LPC lipids as calculated from 9-^2^H choline incorporation 24 hrs after injection by sex and genotype (n=5-9, mean±SEM). For C, * p<.05, ** p<.01, *** p<.001, **** p<.0001 by 2-way unpaired ANOVA, corrected for multiple comparisons by Tukey’s method.

**Supplemental Figure 6:**
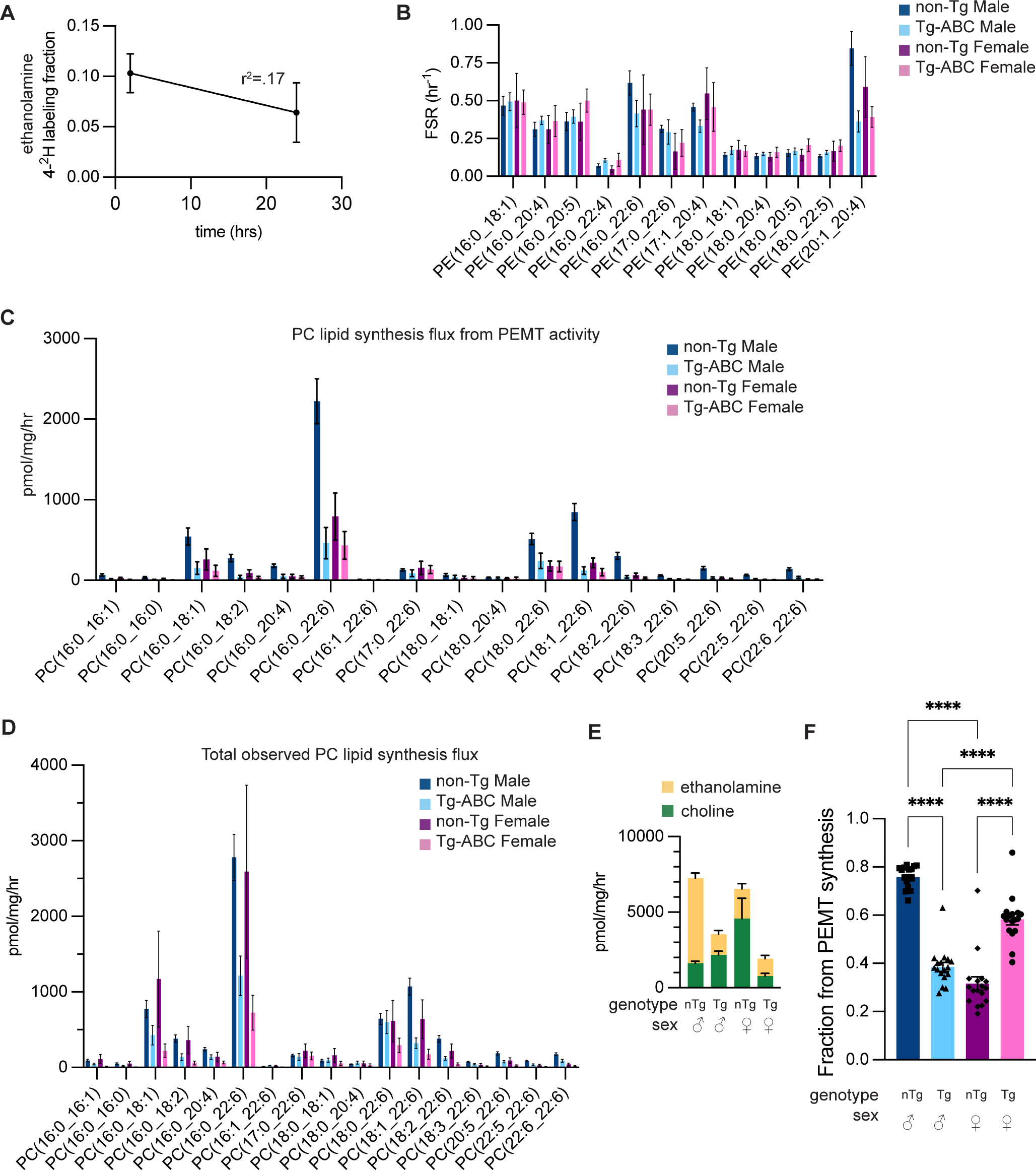
PC lipids synthesis pathways show sex-specific regulation upon activated β-catenin overexpression. A. Recovered labeling fraction of ethanolamine in a combined population of non-Tg and Tg-ABC, male and female zebrafish livers 2 hrs (n=4) and 24 hrs (n=30) after 4-^2^H ethanolamine injection, fit to a linear regression (mean±stdev). B. FSR of PE lipids as calculated from 4-^2^H ethanolamine incorporation into non-Tg and Tg-ABC, male and female zebrafish livers 24 hrs after injection (n=4-13, mean±SEM). C. Absolute synthesis rates of PC lipids via the PEMT pathway as calculated from 4-^2^H ethanolamine incorporation and total lipid abundance 24 hrs after labeled ethanolamine injection by sex and genotype (mean±SEM). D. Absolute synthesis rates of PC lipids via the CHPT1 pathway as calculated from 9-^2^H choline incorporation and total lipid abundance 24 hrs after labeled choline injection by sex and genotype (mean±SEM). E. Total synthesis flux for all PC lipids combined as determined by ethanolamine and choline tracers by sex and genotype (mean±SEM). F. Fraction of total synthesis flux via the PEMT pathway as calculated from labeled ethanolamine incorporation for all PC lipid species by sex and genotype (n=17, mean±SEM). For F, * p<.05, ** p<.01, *** p<.001, **** p<.0001 by 2-way unpaired ANOVA, corrected for multiple comparisons by Tukey’s method.

